# On the Origin of Feces: Fungal diversity, distribution, and conservation implications from feces of small mammals

**DOI:** 10.1101/2021.09.23.460834

**Authors:** Alexander J Bradshaw, Kendra Autumn, Eric Rickart, Bryn T.M. Dentinger

## Abstract

Fungi are extremely diverse, but only a small fraction of the total estimated species have been characterized. Often, the extent of diversity and distribution of fungal communities is difficult or near impossible to assess. This is due to the fact that many Fungi are cryptic and persist predominantly hidden within substrates such as soil or plant material. This is particularly true for hypogeous sporocarps, including truffles, which are extremely difficult to survey in a systematic manner. However, fungi with hypogeous sporocarps have evolved traits that make them highly attractive to animals, such as small mammals, which ingest and disperse fungal spores through defecation. Here, samples of feces from 138 small mammals collected in the western United States were assessed for total fungal diversity using a dual-index metabarcoding, high-throughput Illumina sequencing approach. Our findings exhibit many identifications within *Agaricomycetidae*, with 65 of the 138 samples containing sequences belonging to several species of the hypogeous truffle genus *Rhizopogon*. Metadata, such as geospatial coordinates, for each captured small mammal can be used as a proxy for the presence or absence of *Rhizopogon* species identified in their feces. Utilizing these proxy data, along with publicly available data on observation and occurrence from data repositories such as GBIF and MycoPortal, it is possible to augment our current knowledge of the extent of occurrence and area of occupancy of cryptic hypogeous fungi without direct observation, further enhancing our ability to assess their conservation status.

## Introduction

Kingdom *Fungi* is extremely diverse and ubiquitous across the globe, with ∼148,000 currently accepted species known from the estimated 2.2-12 million distributed on all continents and in most aquatic habitats (Hawksworth and Lücking 2017; Wu et al. 2019). Documentation of fungal diversity has lagged far behind other groups of multicellular organisms, in part due to their generally cryptic habits, their unpredictable production of ephemeral macroscopic sporocarps, and a far greater emphasis on plants and animals that has severely biased biodiversity knowledge (Troudet et al. 2017). Recent technological advances such as molecular identification with high-throughput sequencing have made it possible to document fungal diversity indirectly from environmental samples, greatly accelerating the rate at which fungi can be detected and identified (Chase and Fay 2009; Begerow et al. 2010; Schmidt et al. 2013; Nilsson et al. 2019) . Despite the increasing efficiency in documenting fungal diversity, our knowledge of fungal distributions is still overwhelmingly incomplete. This may be due, in part, to the inefficiency inherent in environmental sampling: small volumes over large areas require large numbers of samples, which is compounded by the generally low biomass of target organisms in each sample. Unlike plant and animal surveys, where the target organisms are more easily observed and more consistently observable over long periods of time or can be baited and trapped, fungal surveys have suffered due to the cryptic nature of their subjects. Only a fraction of fungi produce macroscopic reproductive structures, and even these large sporocarps are often unpredictable in occurrence and appear ephemerally. Therefore, fungal surveys have severely lagged behind plant and animal surveys, with consequences in important areas like taxonomy and conservation. Coupled with the high diversity of fungi, low sampling efficiency creates a double-edged sword that has lead to the vast majority of species being undocumented, undescribed, or known only from DNA sequences (dubbed “dark matter fungi”) (Ryberg and Nilsson 2018).

Many fungi participate in specialized associations with other organisms, especially plants. Therefore, accurate documentation of these fungi is important to understand how these associations function at varying scales, from one-to-one interactions, up to whole ecosystems. For example, ectomycorrhizal fungi that form symbiotic relationships with plant roots can provide important benefits to their host plant, including increased uptake of micronutrients, and receive carbon via photosynthates in a mutualistic exchange (Smith and Read 2010; Courty et al. 2010). This symbiosis is globally important, with an estimated 60% of all woody tree stems belonging to obligately ectomycorrhizal plants (GFBI consortium et al. 2019). Baseline documentation of both partners in this important symbiosis is fundamental to fully understanding it and to accurately predict its response to future change. For example, the presence/absence of host generalist or specialist symbionts, and the relative role they play in maintaining and shaping the symbiosis, is key to understanding its resilience to environmental change (Wilson et al. 2012; Liao, Chen, and Vilgalys 2016).

The IUCN Red List Categories and Criteria provide guidelines based on aspects of a species such as population size, the extent of occurrence, fragmentation, and ongoing changes to these characteristics to place species in one of several categories indicating conservation needs (Endangered, Vulnerable, etc.) (Mace et al. 2008; Rodríguez et al. 2015). These guidelines are designed to be widely applicable to many different types of organisms, including fungi (Cannon et al. 2018). The ability to accurately assess whether an organism can be categorized as threatened or endangered is extremely important within the context of conservation. Inaccurate assessment may lead to the misallocation of resources with respect to the actual risks facing wild populations. Due to the difficulty of locating most fungi, we know relatively little about their distributions, making it difficult to assess whether a particular fungus should receive conservation focus, or if our time and resources are better spent elsewhere.

One approach to improve sampling efficiency for fungi is to use a more reliably sampled source that can function as a proxy for them. For instance, EJH Corner famously trained monkeys to collect fruit samples from the tops of trees at the Singapore Botanic Gardens (Mabberley 1999). Similarly, truffle hunters utilize pigs or trained dogs to locate and excavate prized and commercially valuable culinary delicacies, such as *Tuber magnatum* (the Italian white truffle) often hidden belowground (Riccioni et al., 2016). Like pigs and trained dogs, many small mammals of northern temperate forests, paralleled by marsupials of Australian forests, seek out and consume fungi, mostly truffles and false truffles, sometimes as the majority of their diets (Maser, Trappe, and Nussbaum 1978; Lehmkuhl et al. 2004; Vernes, Cooper, and Green 2015). Truffle-forming fungi have likely been selected to encourage the discovery and consumption of their sporocarps by animals as a means to spread their spores aboveground, necessary due to the loss of ballistospory during the adoption of a hypogeous, sequestrate lifestyle (Johnson 1996). Moreover, because identifications of fleshy macrofungi in small mammal feces almost certainly reflect the presence of sporocarps, due to their small home range and rapid digestions (Padmanabhan et al. 2013; Langer 2002; Hawes 1977), observing fungi in feces are reliable indicators of a species presence at a given location at a given time. This is important, because, unlike environmental DNA, these data points represent reproductively mature individuals that are established and growing, features that are important for assessing species distributions but are inherently lacking in eDNA samples. Fungivorous animals, therefore, are potential proxies for the fungi they consume. They are attracted to scents produced by hypogeous fungi (Splivallo et al. 2011; Stephens et al. 2017) and have the capacity to continuously survey for them. As much as we may envy these abilities, it’s undeniable that humans are at a major disadvantage when it comes to locating wild hypogeous fungi. However, with the use of passive trapping, small mammal feces may represent a source of highly efficient sampling for some fungi, especially truffles and false truffles.

Previous studies have used DNA sequencing of feces to monitor the population of threatened and elusive animals such as snow leopards and grizzly bears, and more recently to identify populations of African elephants which aided in identifying illegal Ivory trading routes. (Janečka et al. 2008; Wasser et al. 2004; 2015; Phoebus et al. 2020). Fecal sampling has also been used to identify the contents of animal diets, such as in rodents, hares, and even river otters (Cloutier et al. 2019; Buglione et al. 2018; Elliott et al. 2020; Harper et al. 2020), but focused primarily on community identification, and in the context of the host diet rather than the identity of community members. These studies show how powerful and informationally rich fecal sampling can be for population distribution studies While these previous studies focused on macrofauna, this same approach could be used for consumed fungi, which are often used as food sources for many metazoan taxa ranging from fungivorous arthropods to opportunistic small mammals and even many great apes (Yamashita et al. 2015; Pyare and Longland 2001; Hanson, Hodge, and Porter 2003, Elliott et al. 2020). In this study, we present the idea that the known ecology of a host along with sample metadata, such as the geospatial coordinates, can be informative and used as a proxy for a rough estimation of community member distribution in the case of difficult to find macrofungi. To meet this objective, we set out to address four questions, **1) Can DNA metabarcoding of fecal samples be used to identify fungi associated with small mammal consumption, such as those with hypogeous sporocarps? 2)What diversity of fungus species can be recovered in a diverse collection of small mammal feces, and is there any distinct host diet specialization among them? 3) Can metadata, such as the geographic location of trapping, extend our understanding of the diversity, ecology, and distribution of fungi.? 4) Do these “collections” aid in making meaningful conclusions about the distribution of fungal species that can be used to inform possible conservation efforts?**

## Results

### Raw sequencing data and ASV information

After the quality processing step of DADA2, and removing all singleton ASVs of our 138 fecal samples, 136 samples were left for further analysis. Read and variant processing with DADA2 (Callahan et al. 2016) along with taxonomic identification based on the UNITE fungal ITS database, found 4650 unique ASV’s, representing 531 genera across all samples. Our ASVs were dominated primarily by fungi belonging to the phyla *Ascomycota* and *Basidiomycota*, with the genera *Mycosphaerella* and *Rhizopogon* being the two most abundantly found ASVs (figure 1). Additionally, other fungal orders of note found in high quantity were *Mucorales* (dung dwelling fungi), *Pleosporales* (common saprobic fungi found on decaying plants), *Agaricales* (Fungi producing Mushroom macro reproductive structures), *Tremellales* (dimorphic fungi which can produce gelatinous macro reproductive structures), and *Pezizales* (fungi which include economically important fungi such as morels and black and white truffles). Due to its known hypogeous morphology, generally common distribution, and its large presence within our sample set, the genus *Rhizopogon* was chosen for more in-depth analysis. Subsetting the dataset for only members of the *Rhizopogon* genus yielded 65 samples, which contained 580 total ASVs. Of these 580, 394 were assignable to species level, representing 11 *Rhizopogon* spp., while 186 could only be identified to genus level. Assignable ASVs were collapsed based on species assignment, and unassignable ASVs were clustered together at 99% similarity, yielding 11 assignable taxa, and 11 Operational Taxonomic Units (OTUs) for further analysis.

**Figure 1:**
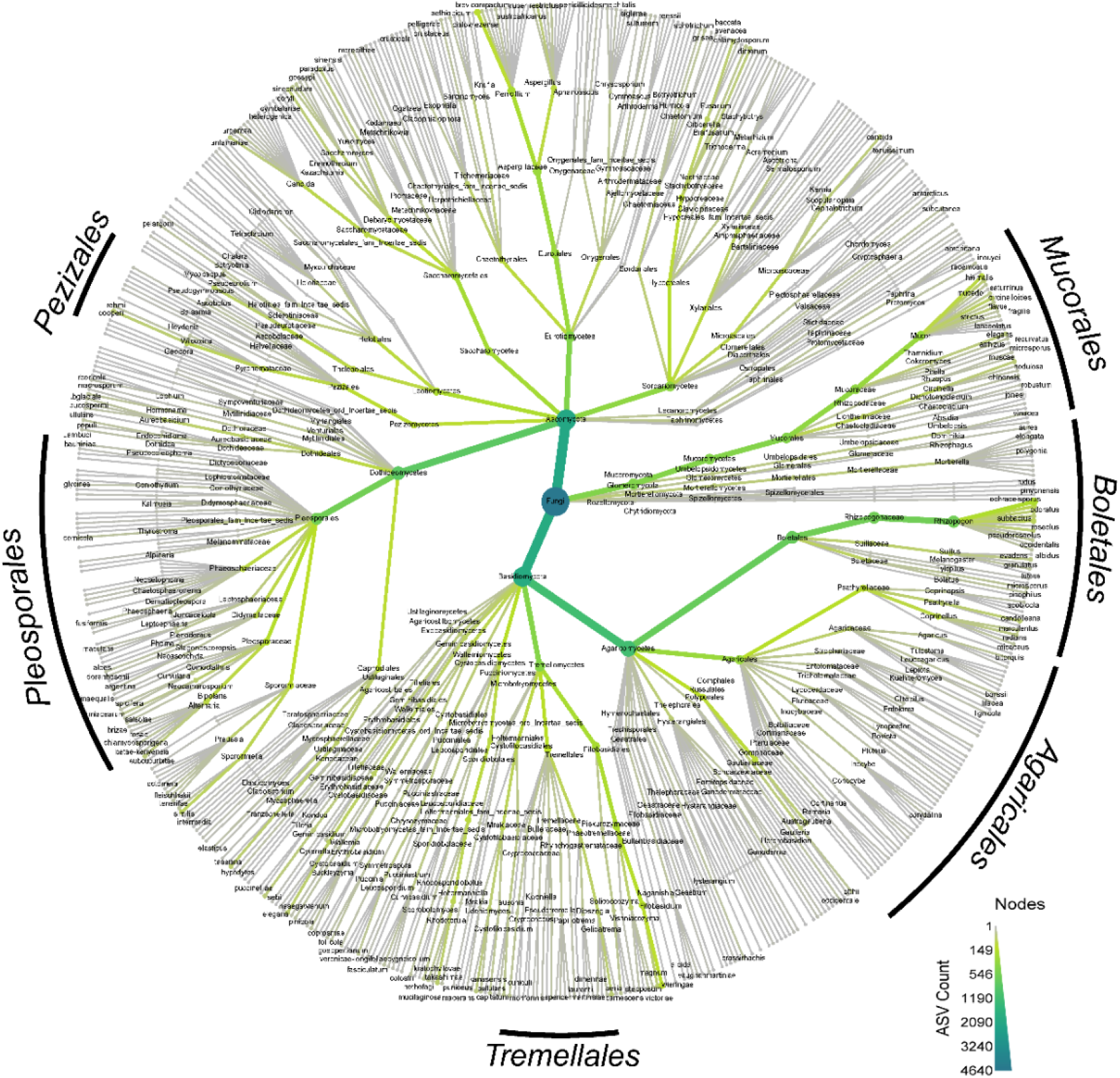
Leaflet composition of Total Fungal Community. Leaflet distribution of the full fungal community across all samples, with a large number of ASVs are indicated with thicker lines as well as darker coloration. Major orders are labeled for quick reference.

### Phylogenetic analysis of recovered N/A *Rhizopogon* distribution

With 186 of our *Rhizopogon* ASVs unassignable to species-level taxonomy, we chose to utilize phylogenetic analysis to further investigate *Rhizopogon* diversity. To do this we parsed the general release UNITE database (version 8.2, 04 April 2020) (Abarenkov et al. 2010) for all species hypothesis sequences from the family *Rhizopogonaceae* followed by ITSx (Bengtsson-Palme et al. 2013) to extract the ITS2 sequence for each entry to standardize our amplicons along with the UNITE “species hypothesis” sequences (Nilsson et al. 2014). We then generated a phylogenetic tree, containing all UNITE sequences, assigned *Rhizopogon* ASVs collapsed to species level, and our unassigned ASVs clustered into OTU’s at 99%. Unfortunately, NA OTUs were not able to be resolved using phylogenetic analysis, most likely due to a lack of diversity representation found within the reference database.

We wanted to further investigate and compare *Rhizopogon* diversity across our mammal species. To do this we calculated Faith’s Phylogenetic Diversity (PD), a measurement of biodiversity based on phylogenetic analysis for each mammal species (Faith 2018). PD of our *Rhizopogon* consuming small mammals ranged from PD= 1.262573 belonging to the taxa *Peromyscus maniculatus,* to PD=0.213792 belonging to the taxa *Dipodomys merriami* (Figure 2). This would suggest that some small mammal species are consuming a wider diversity of truffles compared to others. However, when each sample is investigated separately, PD values fluctuate from sample to sample within a single small mammal species (Supplementary Table 1). We looked into this further using our highest PD *Peromyscus maniculatus* samples and found that all 4 of them had been collected on the same day and within the same geolocation. However, one sample, UMNH.Mamm.43034, corresponds to a high PD value of 0.775310717 while the others corresponded to values between 0.194667809 and 0.282245014 with little overlap in *Rhizopogon* taxa. This suggests that individual small mammal species are less important than geographic locality, and consumption is likely to be generalist and opportunistic in nature, a characteristic that has been studied and is considered to be a vital role in forest ecology (Stephens and Rowe 2020).

**Figure 2:**
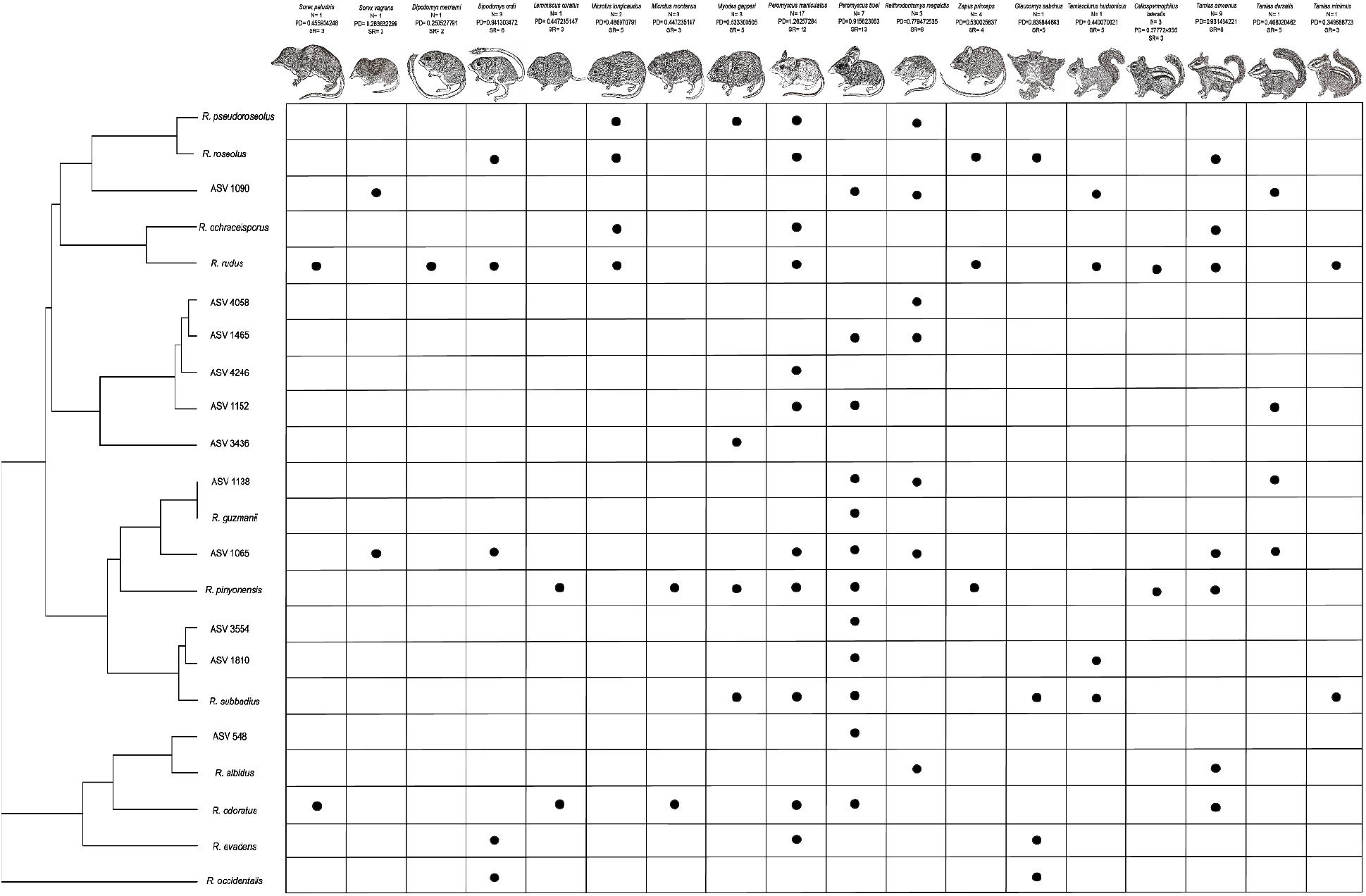
*Rhizopogon* ITS2 phylogenetic tree and presence/absence in each small mammal species along with reported Faith’s phylogenetic diversity and Species richness found in each. (Left) Phylogenetic cladogram of *Rhizopogon* ASVs and OTUs corresponding to the presence or absence in each small mammal taxa. (Top) All small mammal taxa as well as calculated faith’s phylogenetic diversity, species richness in dataset as well as number of collected specimens.

### Environmental factors associated with *Rhizopogon* presence

Alpha diversity measurements of *Rhizopogon* in our samples were conducted using both Choa1 and Shannon indices (Supplementary Figure 4). Alpha diversity across our samples did not indicate that any single species of small mammal was associated with *Rhizopogon* consumption diversity. However, using metadata and the geospatial coordinates for each sample trapping occurrence, we were able to retrieve bioclimatic variables (Fick and Hijmans 2017) and perform a principal component analysis for both qualitative and quantitative variables. In regard to sample variables, we found no discernable correlation between small mammal sex (supplementary figure 3) and the presence of *Rhizopogon* ASVs. This suggests that in our sample set one sex is not more likely to consume *Rhizopogon* than the other. However, our data did find structure related to the season in which they were collected (Figure 3a). We further dissected the bioclimatic data to see which variables contributed most to the community within our samples. We found that 77.38 % of the variability within our *Rhizopogon* consumption set was due to two dimensions (Figure 3b). Within those dimensions we found the highest correlation to be associated with precipitation and warm temperature patterns which match with the observation of the individual PCA having an association with seasonality, which has been studied before in *Rhizopogon* (Luoma, Frenkel, and Trappe 1991; Hunt and Trappe 1987).

**Figure 3:**
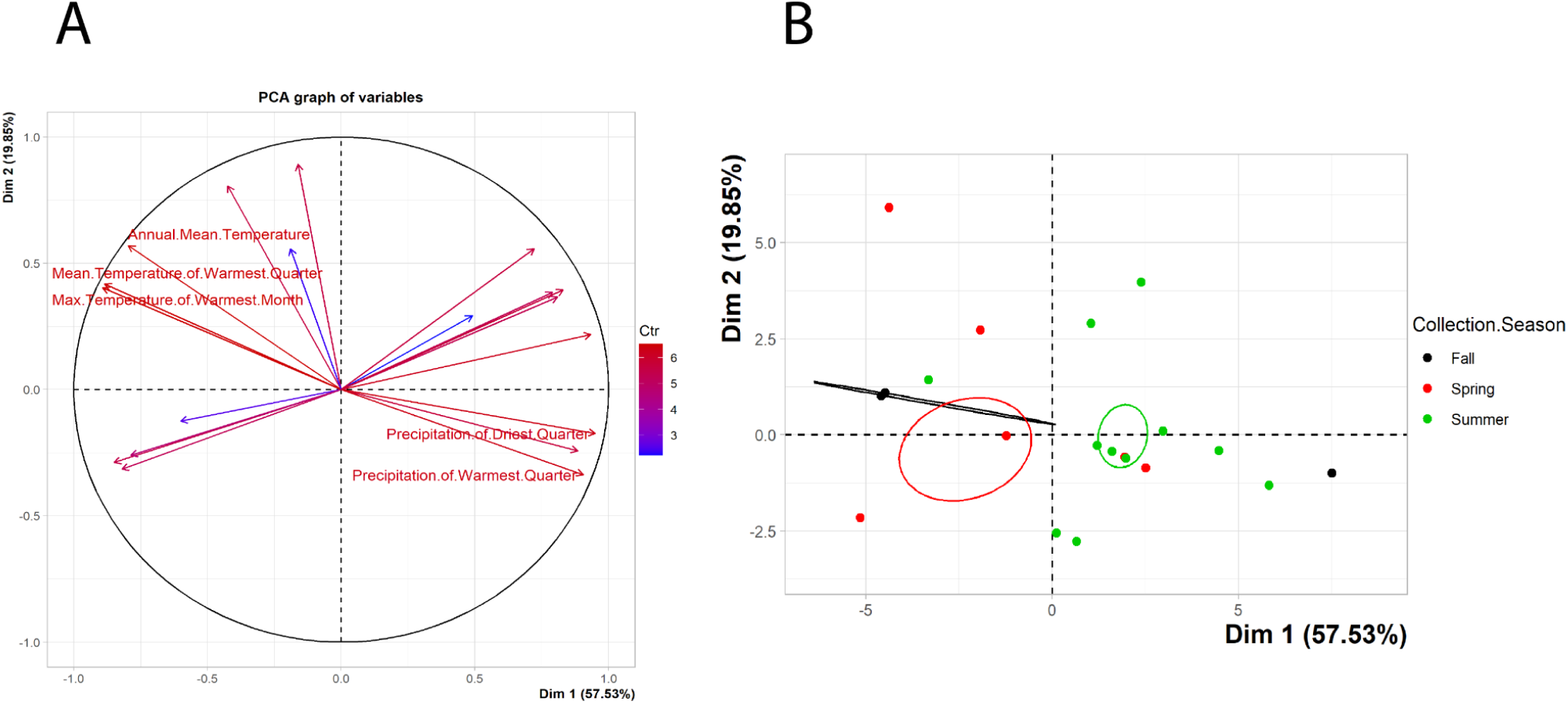
Principal Component analysis of the top 5 Bioclim variables associated with *Rhizopogon* presence in samples and confidence ellipse in seasonality. (Left, A) Principal component analysis of sample bioclimatic variables. More Contribution to variability is shown in red with the top 5 variables being labeled. (Right, B) Principal component analysis of samples based on season of collection with confidence ellipses.

### Geocat EOO and AOO

The extent of occurrence (EOO) and area of occupancy (AOO) are two criteria that are commonly used for IUCN risk designation (Rodríguez et al. 2011; Schatz 2009; Rodríguez et al. 2015). As such, they are important measurements to apply to our dataset to determine if *Rhizopogon* fits within a threatened species threshold. Expansion of known ranges of *Rhizopogon* using small mammal trapping location as a proxy data for the known EOO and area of occupancy AOO were generated by importing observation and collection geopoints from GBIF and Mycoportal (Miller and Bates 2017)into Geocat, a tool supported by IUCN for the calculation of EOO and AOO for this purpose (Bachman et al. 2011). Coordinates of trapped small mammals were counted as a single “observation” if an ASV within that sample could be taxonomically identified to species level. Geocat’s mapping feature allowed for graphical representation of EOO and AOO as well as assigning an IUCN threatened category for each species. Across our assigned *Rhizopogon* taxa, *Rhizopogon guzmanni* had the fewest publicly available data points (4), while *Rhizopogon roseolus* had the most data points (900+). The discrepancy in data points for *Rhizopogon roseolus* is most likely due to its commercial success as a cultivable symbiont that is often used in the reforestation and plantation of pine trees (Dunstan, Dell, and Malajczuk 1998; Sousa et al. 2011). Calculations of EOO and AOO for all identified *Rhizopogon* species for each data set as well as those for all data sets combined are reported in Table 2.

**Table 1:**
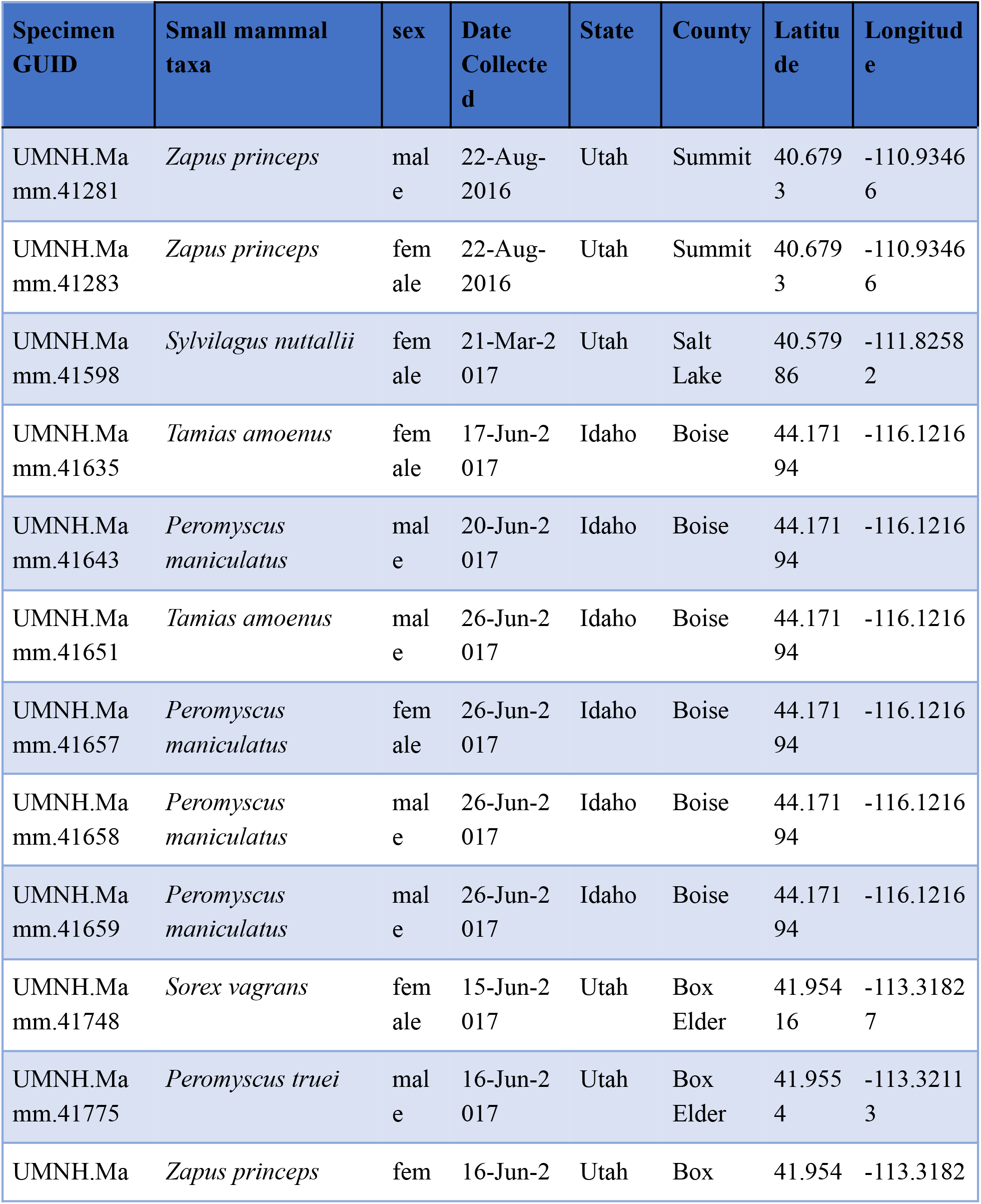
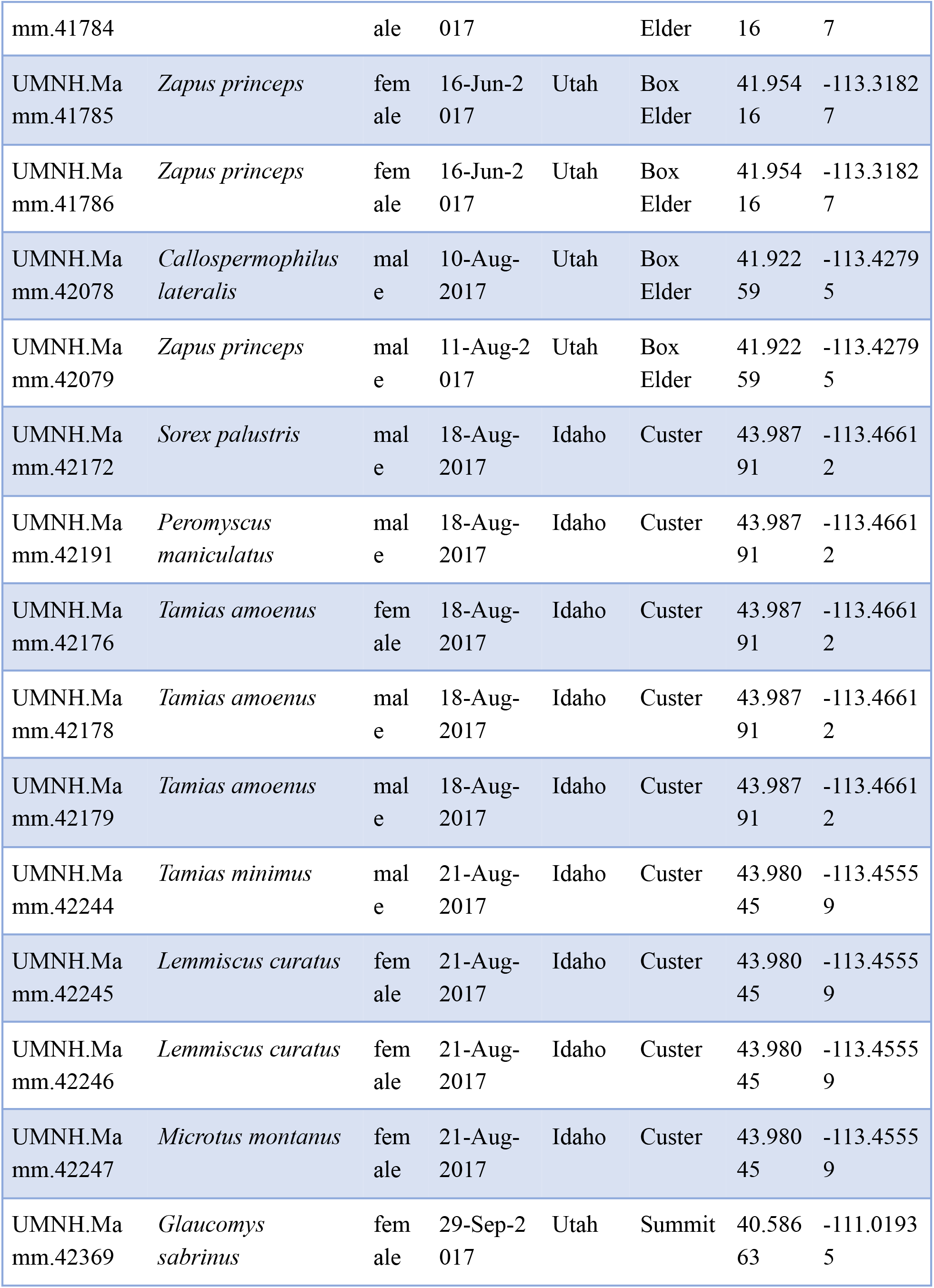
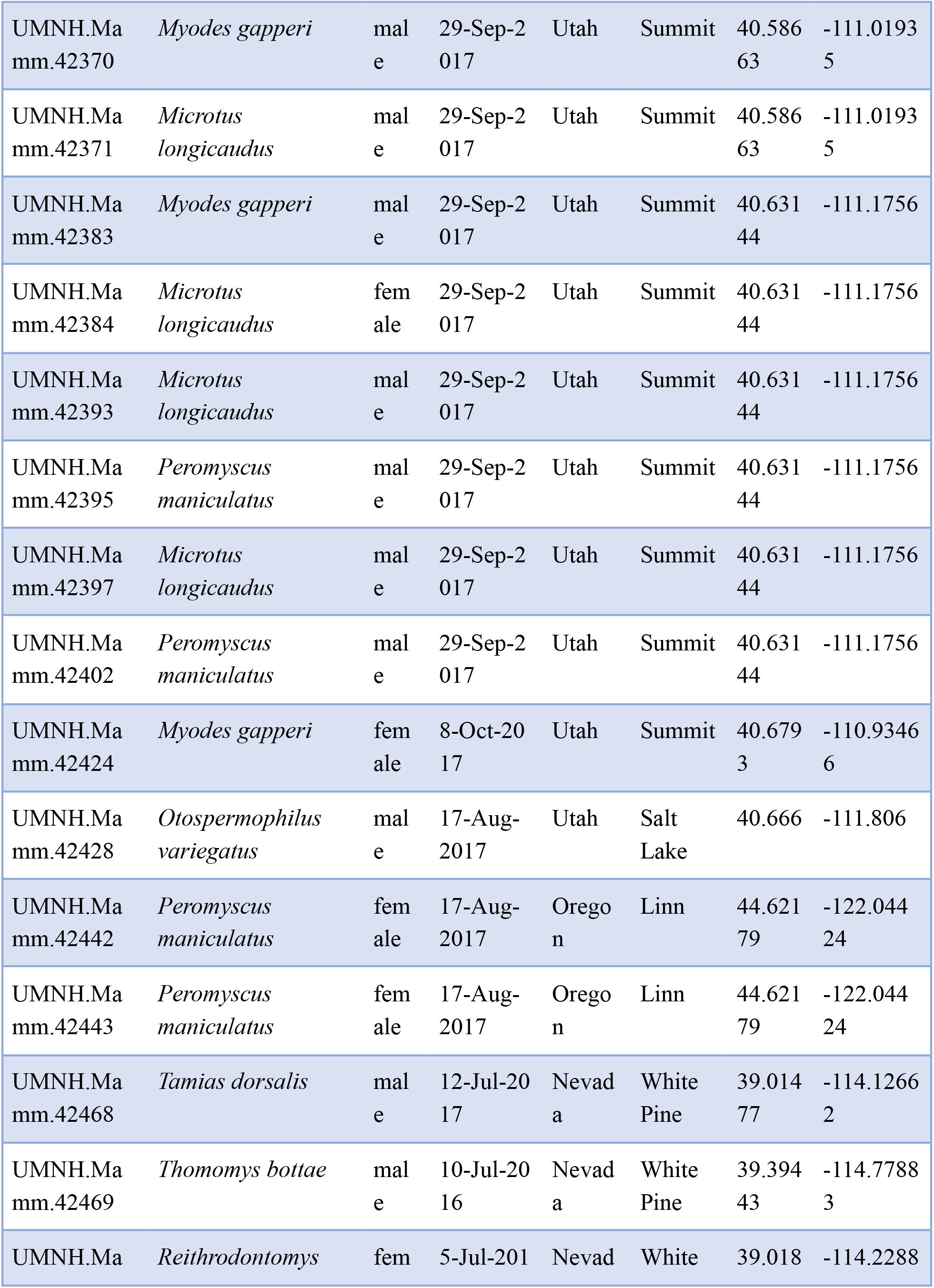
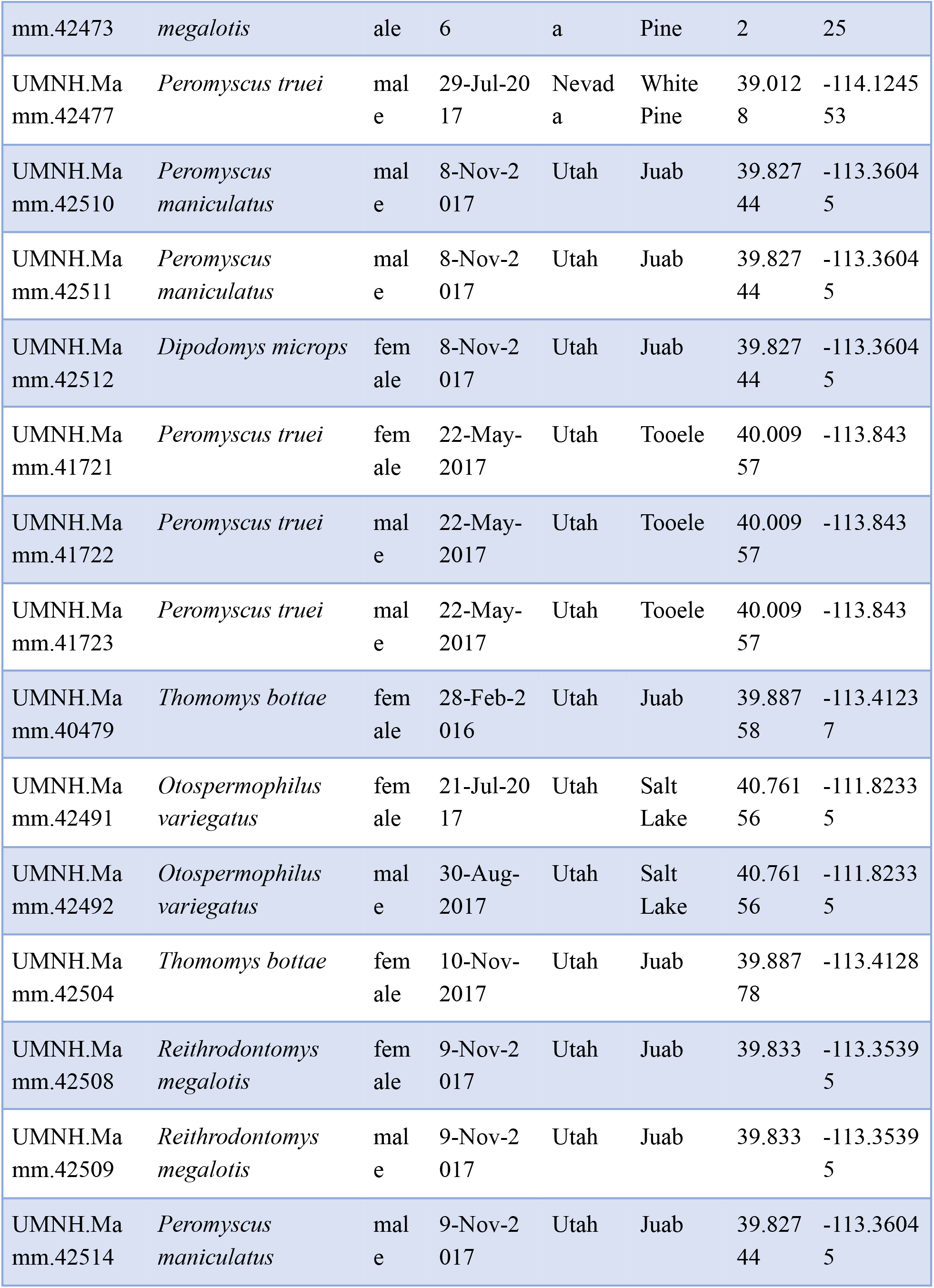
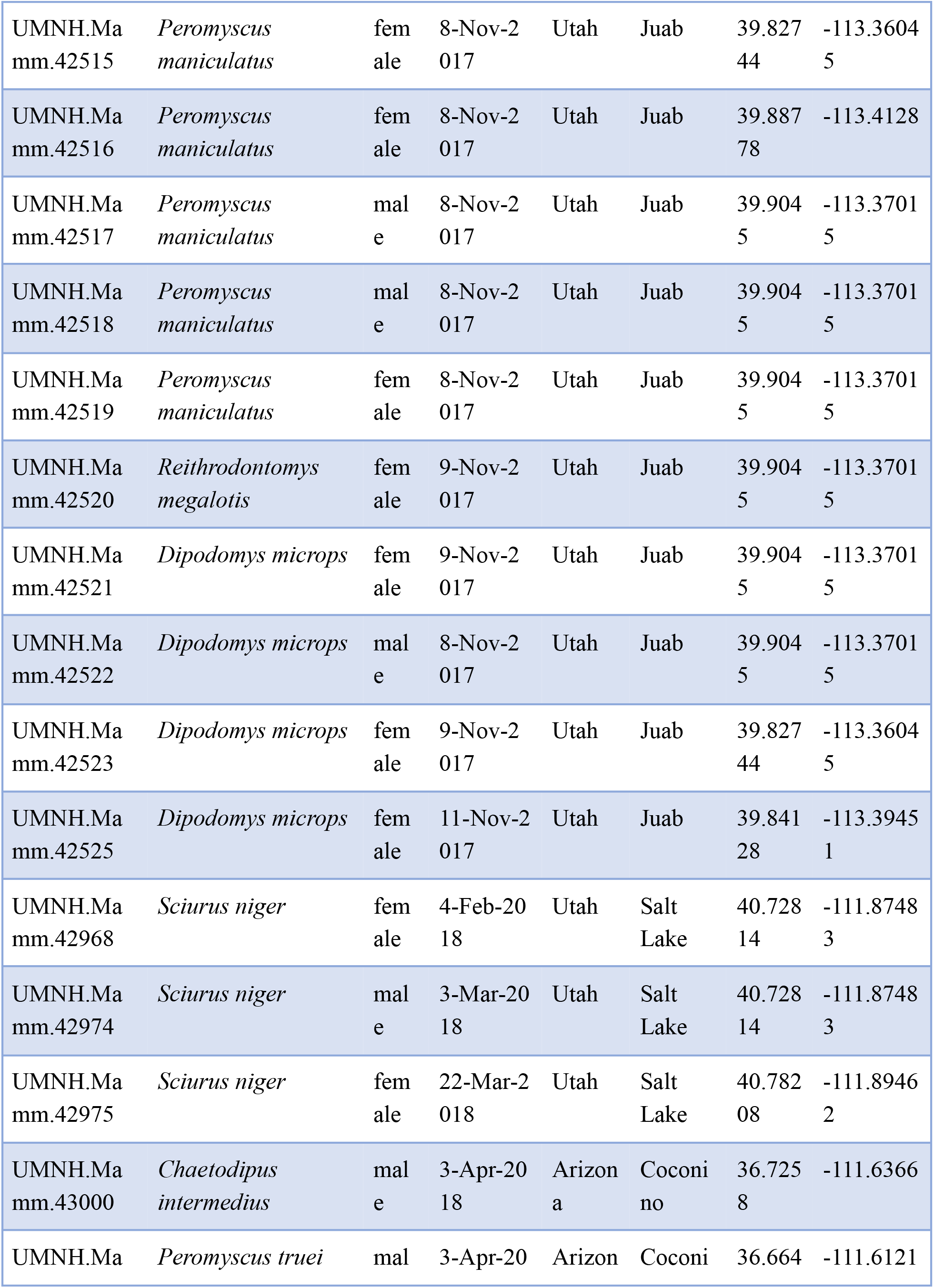
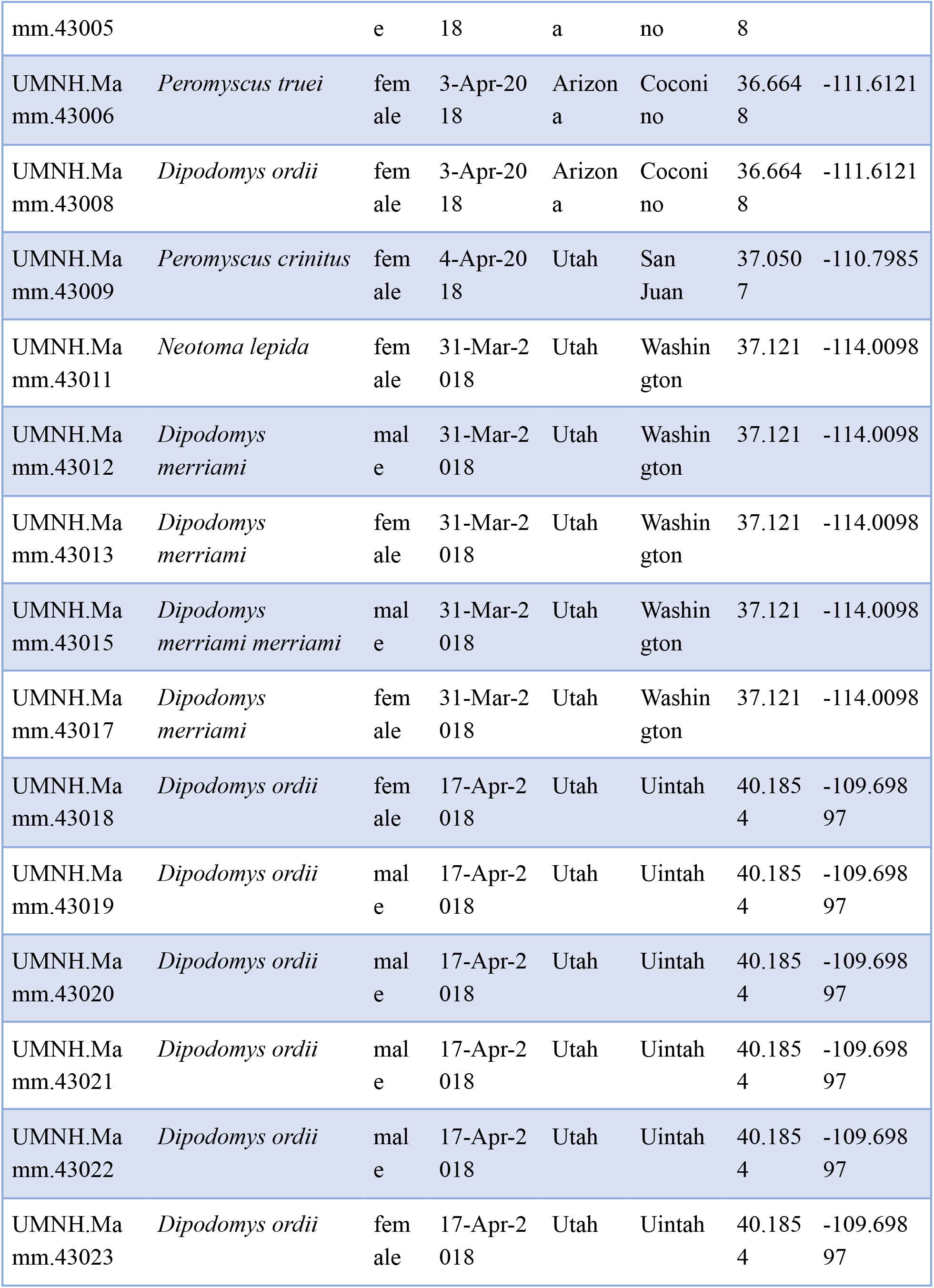
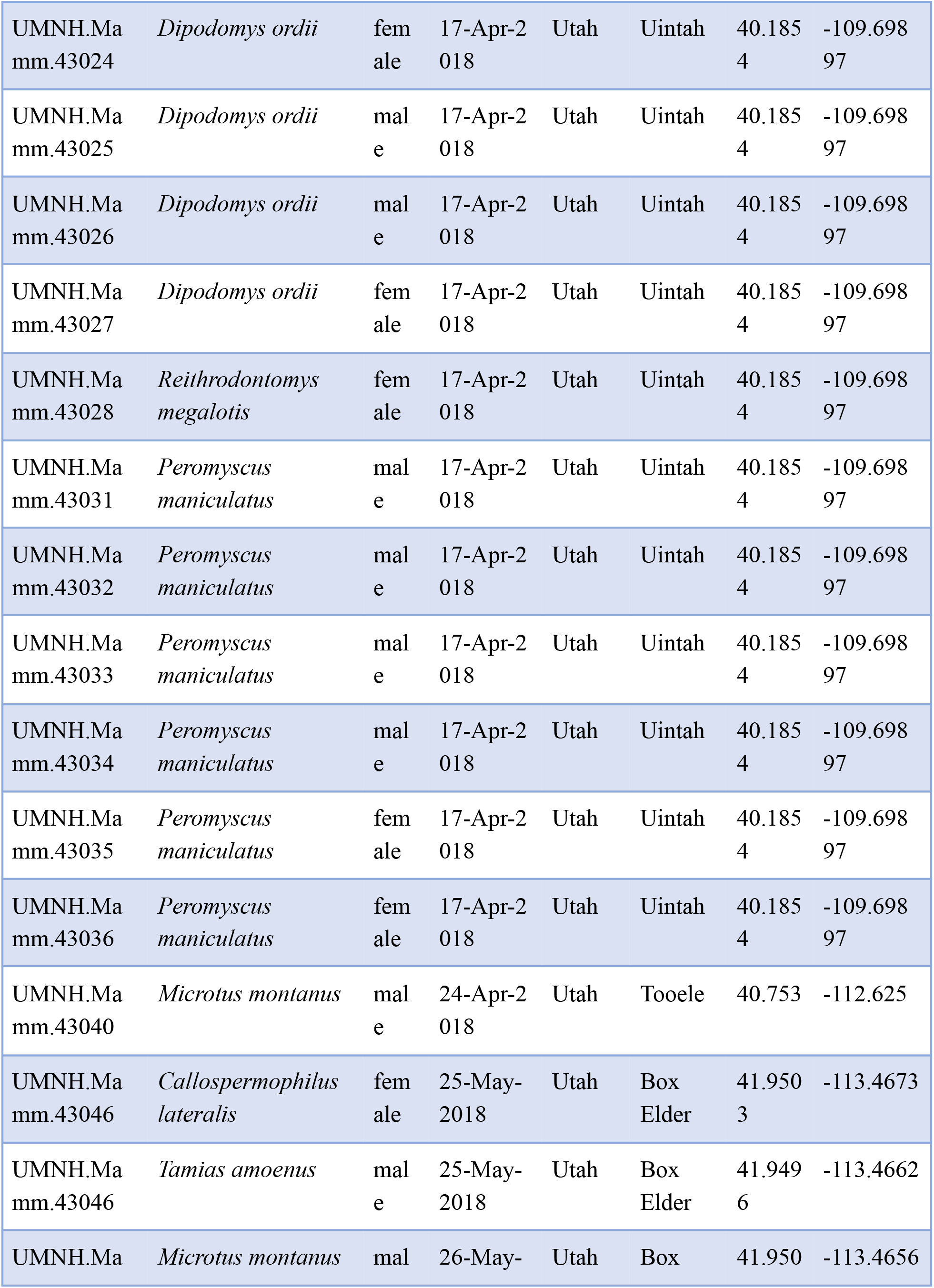
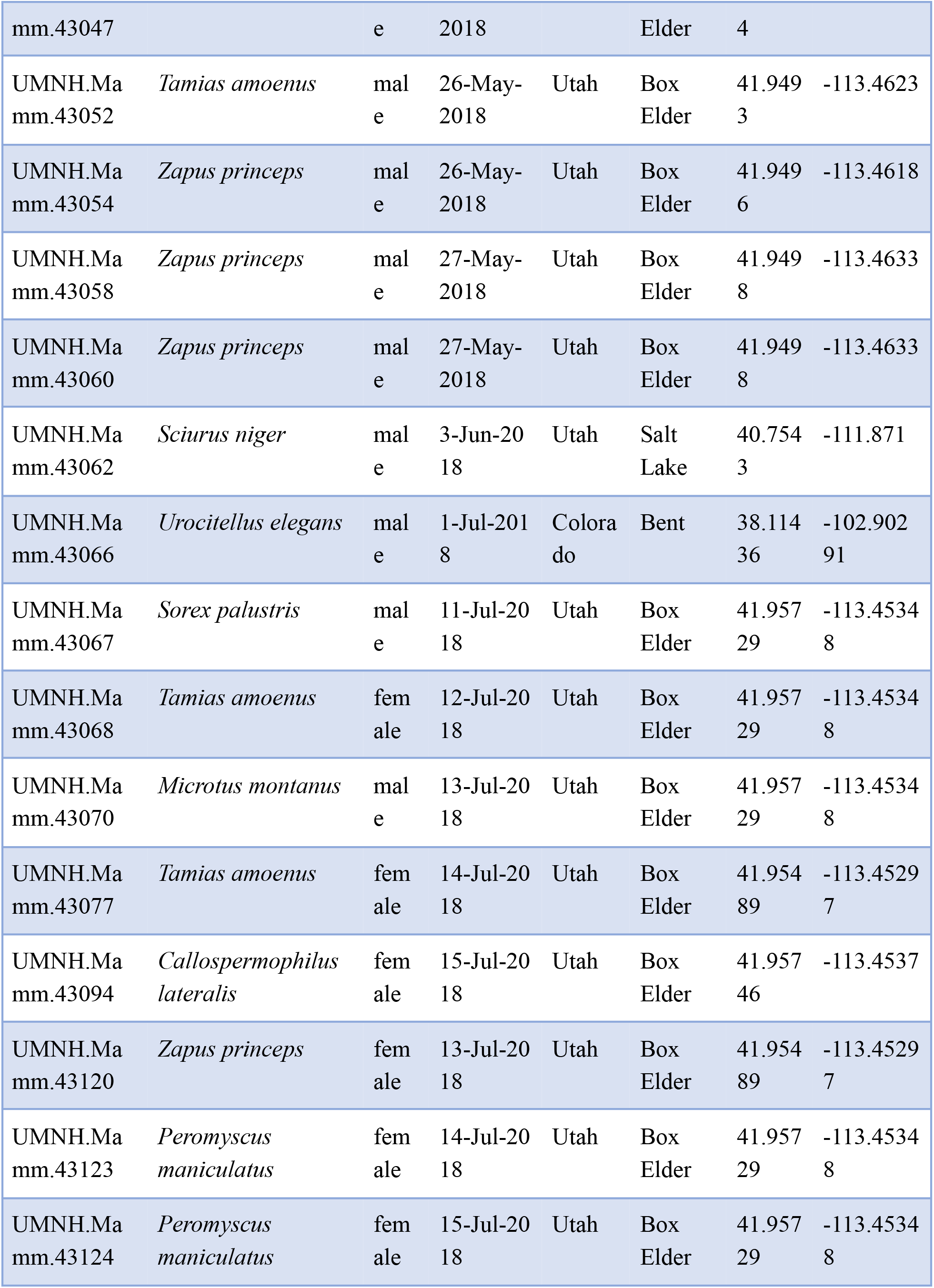
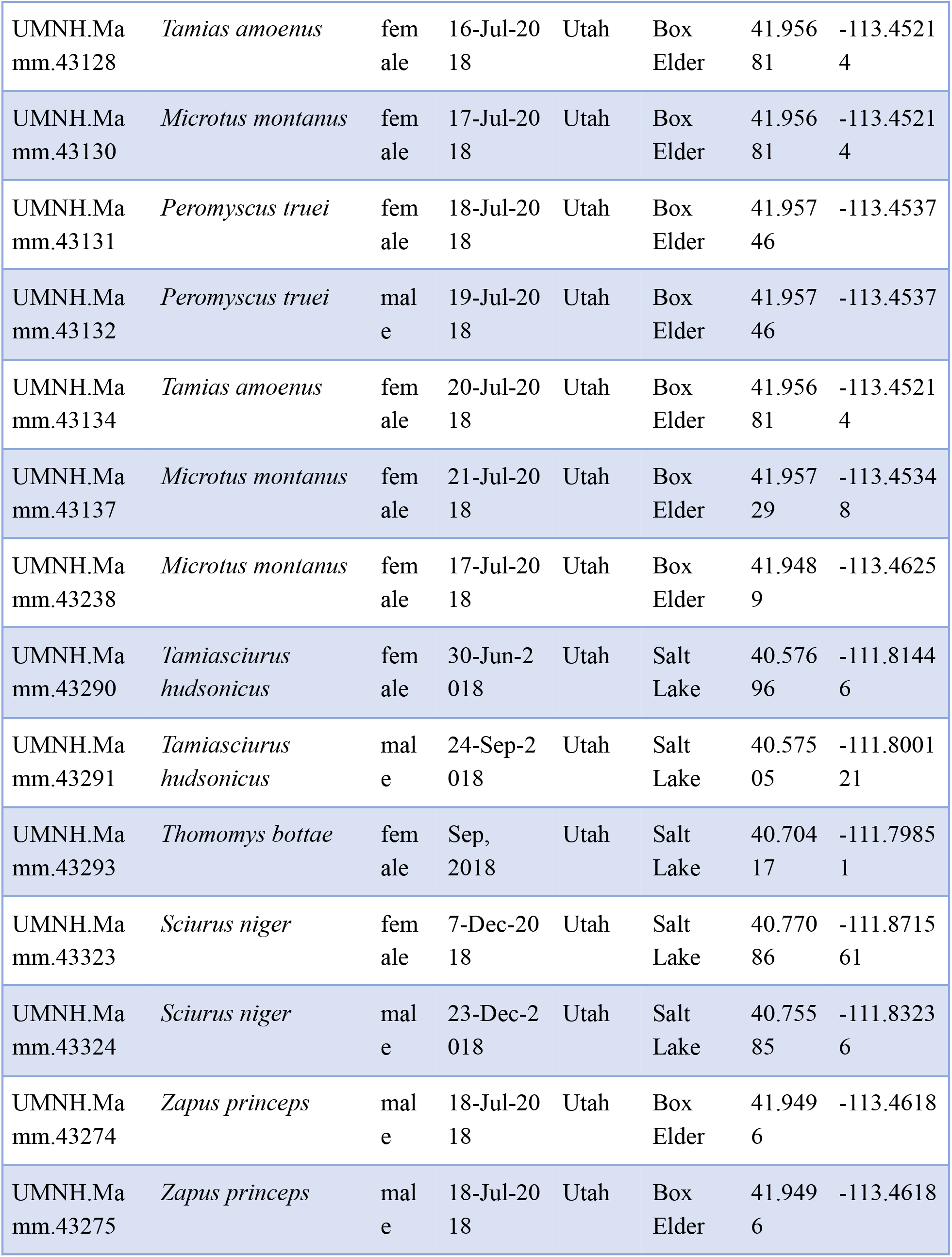
Voucher Table of all specimens used in this study. Table Containing all voucher information for each small mammal including accession numbers, locality, and time of year of each trapping.

**Table 2:**
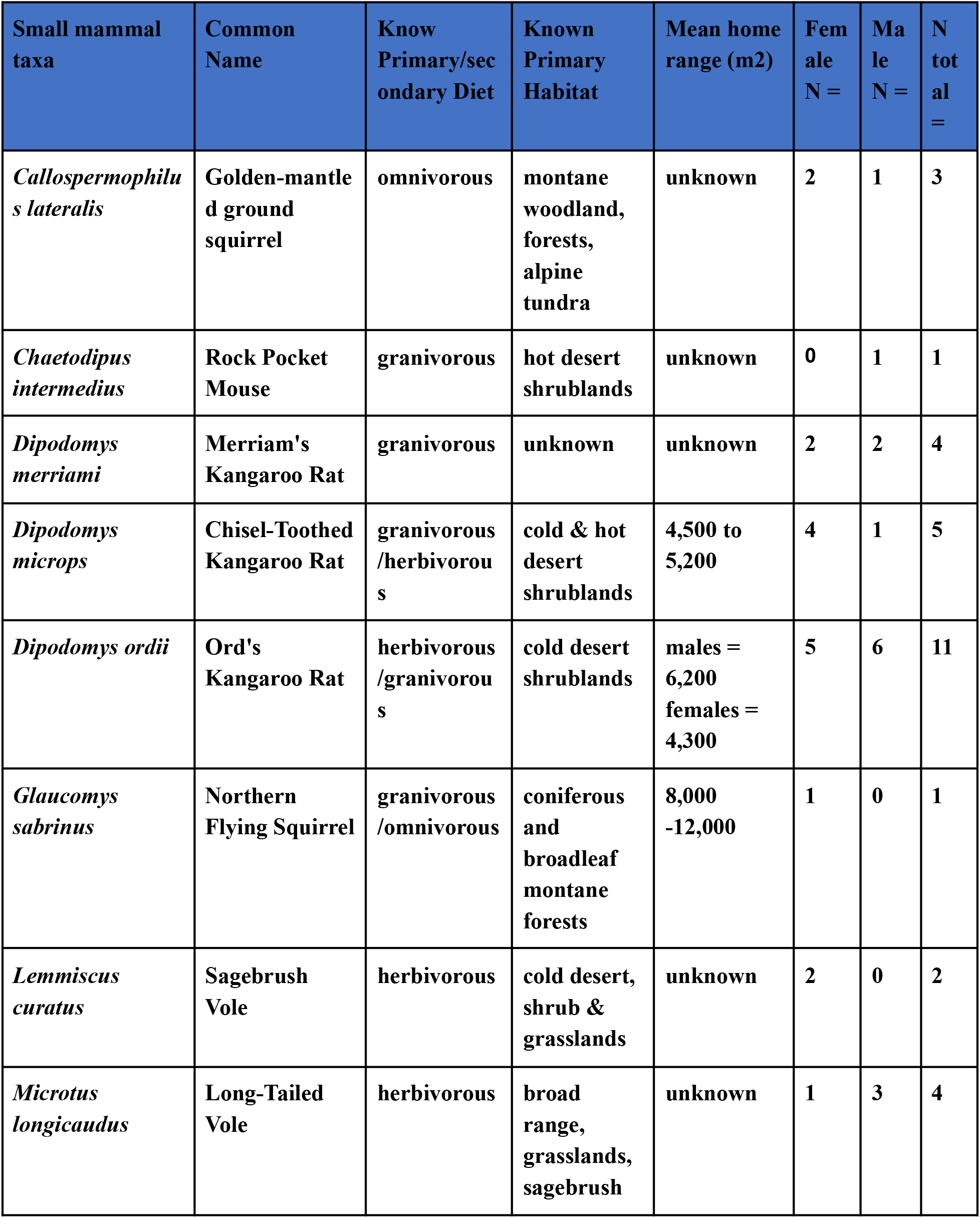
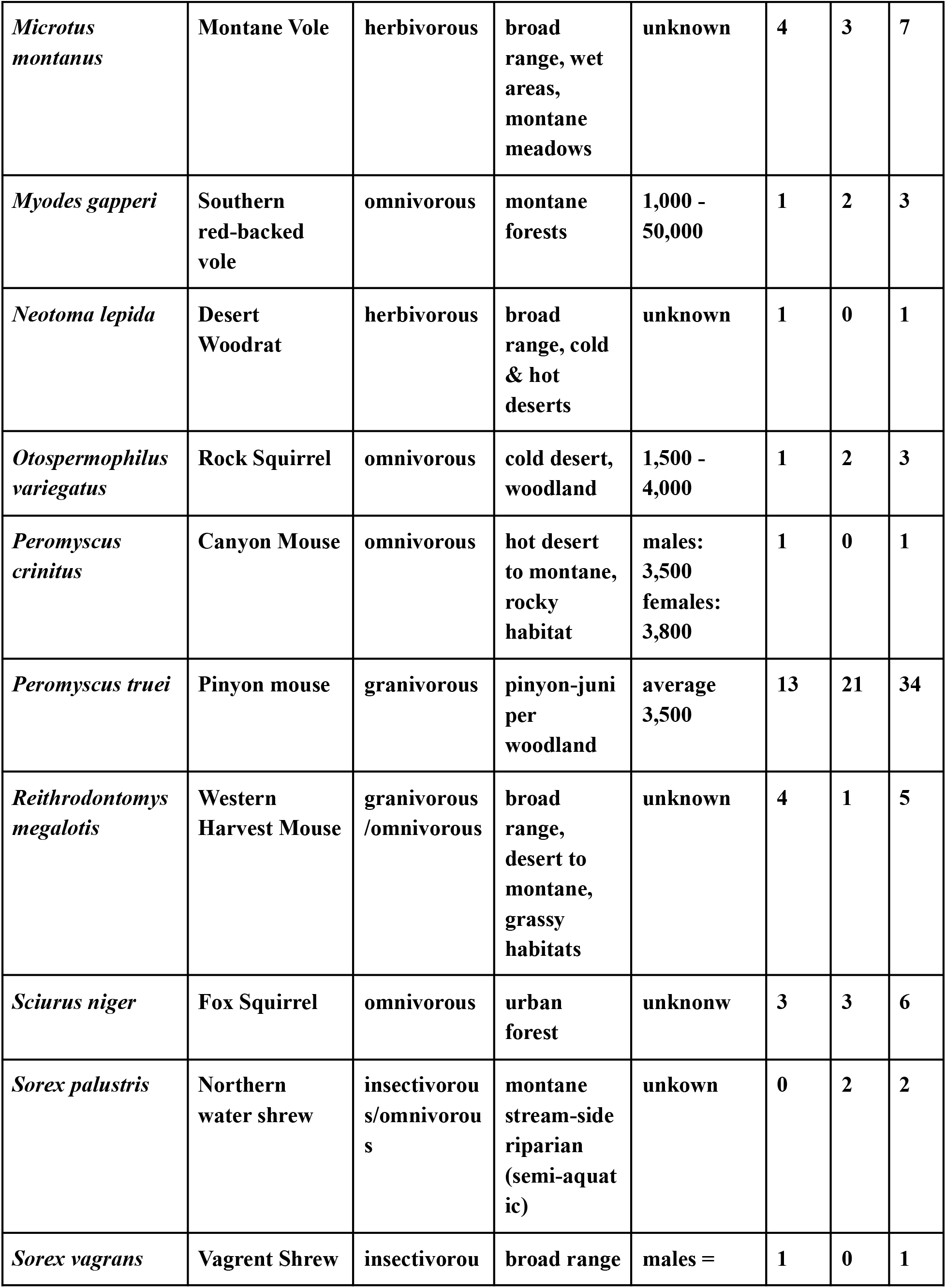
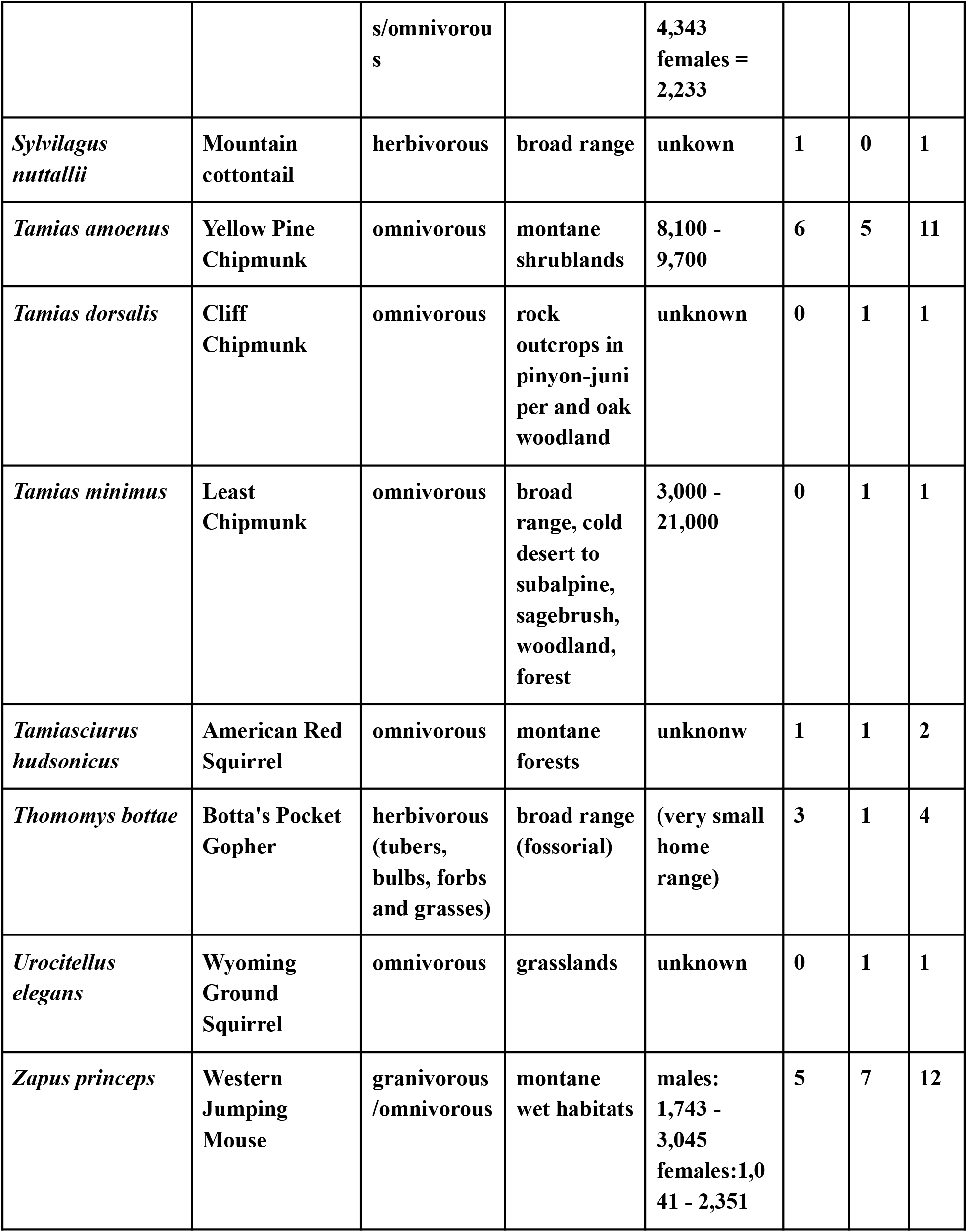
Information table of each small mammal species. Ecological and Behavioral information for each small mammals including home range if known

**Table 3:** Expansion of EOO and AOO and red list status. Values calculated for Extent of Occurrence and Area of Occupancy from the public databases GBIF and Mycoportal. Table also includes values calculated for our sampling, and all data sources combined. IUCN status is derived from all data points combined.

|  |  |  |  |  |  |  |  |  |  |  |  |  |  |  |  |
| --- | --- | --- | --- | --- | --- | --- | --- | --- | --- | --- | --- | --- | --- | --- | --- |
| *Rhizopogon albidus* | 1 | 0 | 2 | 9 | 171<br>637<br>.16<br>6 | 12 | 2 | 0 | 8 | 309<br>541<br>.08<br>8 | 20 | 137<br>903.<br>922 | 8 | Leas<br>t<br>conc<br>ern(<br>LC) | End<br>ang<br>ere<br>d(E<br>N) |
| *Rhizopogon evadensis* | 24<br>1 | 71<br>68<br>20<br>4.<br>22<br>1 | 22<br>4 | 30<br>4 | 339<br>247<br>4.2<br>18 | 392 | 3 | 418<br>68.<br>013 | 12 | 831<br>568<br>4.5<br>19 | 436 | 114<br>748<br>0.29<br>8 | 44 | Leas<br>t<br>conc<br>ern(<br>LC) | End<br>ang<br>ere<br>d(E<br>N) |
| *Rhizopogon guzmanii* | 4 | 0 | 4 | 0 | 0 | 0 | 1 | 0 | 0 | 0.3<br>34 | 8 | 0.33<br>4 | 8 | Criti<br>cally<br>End<br>ang<br>ered<br>(CR) | Criti<br>cally<br>End<br>ang<br>ered<br>(C<br>R) |
| *Rhizopogon occidentalis* | 14<br>3 | 20<br>08<br>33<br>26<br>.0<br>7 | 35<br>6 | 23<br>7 | 127<br>285<br>802<br>.2 | 528 | 3 | 256<br>70.<br>671 | 12 | 124<br>702<br>859<br>.8 | 656 | -258<br>294<br>2.40<br>6 | 128 | Leas<br>t<br>conc<br>ern(<br>LC) | Vul<br>ner<br>able<br>(VU<br>) |
| *Rhizopogon ochraceisporus* | 13 | 11<br>62<br>68<br>.0<br>95 | 36 | 30 | 132<br>956<br>72.<br>96 | 96 | 6 | 921<br>53.<br>084 | 16 | 132<br>956<br>72.<br>96 | 112 | 0 | 16 | Leas<br>t<br>conc<br>ern(<br>LC) | End<br>ang<br>ere<br>d(E<br>N) |
| *Rhizopogon odoratus* | 1 | 9 | 4 | 1 | 0 | 4 | 9 | 603<br>86.<br>999 | 24 | 103<br>927<br>.86 | 28 | 435<br>40.8<br>61 | 4 | Leas<br>t<br>conc<br>ern(<br>LC) | End<br>ang<br>ere<br>d(E<br>N) |

|  |  |  |  |  |  |  |  |  |  |  |  |  |  | LC) | N) |
| --- | --- | --- | --- | --- | --- | --- | --- | --- | --- | --- | --- | --- | --- | --- | --- |
| *Rhizopogon pinyonensis* | 2 | 0 | 4 | 4 | 127<br>31.<br>813 | 12 | 14 | 192<br>892<br>.00<br>7 | 36 | 455<br>181<br>.84<br>4 | 52 | 262<br>289.<br>837 | 16 | Leas<br>t<br>conc<br>ern(<br>LC) | End<br>ang<br>ere<br>d(E<br>N) |
| *Rhizopogon pseudo roseolus* | 76 | 97<br>97<br>06<br>07<br>.9<br>7 | 15<br>2 | 19 | 353<br>255<br>25.<br>83 | 44 | 4 | 949<br>0.9<br>24 | 12 | 100<br>174<br>948<br>.9 | 168 | 220<br>434<br>0.92<br>7 | 16 | Leas<br>t<br>conc<br>ern(<br>LC) | End<br>ang<br>ere<br>d(E<br>N) |
| *Rhizopogon roseolus* | 90<br>0 | 16<br>17<br>03<br>89<br>9.<br>1 | 24<br>80 | 66<br>1 | 238<br>,17<br>4,1<br>72.<br>425 | 160<br>4 | 9 | 139<br>838<br>.44 | 32 | 236<br>678<br>150<br>.5 | 353<br>6 | 749<br>742<br>51.4<br>6 | 105<br>6 | Leas<br>t<br>conc<br>ern(<br>LC) | Not<br>Thr<br>eate<br>ned<br>(NT) |
| *Rhizopogon rudus* | 1 | 0 | 4 | 2 | 0 | 8 | 26 | 214<br>266<br>.26<br>5 | 52 | 657<br>744<br>.04<br>8 | 60 | 657<br>744.<br>048 | 52 | Leas<br>t<br>conc<br>ern(<br>LC) | End<br>ang<br>ere<br>d(E<br>N) |
| *Rhizopogon subbadius* | 1 | 0 | 4 | 2 | 0 | 8 | 6 | 408<br>00.<br>009 | 20 | 199<br>068<br>.89<br>9 | 28 | 158<br>268.<br>89 | 8 | Leas<br>t<br>conc<br>ern(<br>LC) | End<br>ang<br>ere<br>d(E<br>N) |

## Discussion

Fungal communities and species distributions are notoriously difficult to study and map, often due to inconspicuous growth forms dominating their lifecycle. Even though Fungi with macroscopic reproductive structures (e.g., mushrooms, sequestrate “truffles”, etc.) have the longest history of ecological study (Gajos Małgorzata and Hilszczańska Dorota 2013; Hao et al. 2020), baseline information such as known locality and species richness for many macrofungi is largely nonexistent due to difficulty in locating sporocarps, inaccurate taxonomic identification, and lack of long-term monitoring projects (van der Linde et al. 2012). Sampling from feces, or other environmental substrates such as soil, can be non-invasive and enable long-term studies that are less burdened by the difficulty of locating sporocarps in the field. Further, massive collections of fecal material (as well as environmental soil samples), such as those gathered by the NEON project (Dalton 2000; Hopkin 2006), already exist and have metadata readily available for each collection. Utilizing established collection resources could expedite the process of extensive sample collection and analysis required to answer larger-scale fungal community-level questions.

Our study began with the simple idea of using readily available fecal samples from small mammals collected routinely for the Natural History Museum of Utah to investigate fungal diversity in the feces of small mammals using techniques generally used to investigate microbial communities. Metabarcoding of environmental samples has become a popular technique to interrogate organismal communities which would otherwise be difficult to study (Compson et al. 2020; Dieleman et al. 2015; McGee et al. 2019; Andújar et al. 2015; Al Ashhab et al. 2021; Kesanakurti et al. 2011). Here we used the same strategy to target fungal communities in small mammal feces from 138 specimens and were able to identify 4,650 ASVs in 531 genera of Fungi. Although our methods may not have recovered the total fungal community due to technical biases such as variation in DNA extraction efficiency across taxa and primer bias during PCR amplification, our data represent a rich assemblage of macro-and microfungi present in the feces of small mammals. While a mock community of fungi may help assess the degree of community recovery in studies like this, there is currently no mock community that would be representative of the fungi we detected in our samples. Moreover, the added complexity introduced by extracting from fecal material would be difficult to replicate in a mock community without, e.g., feeding sporocarp tissue to small mammals and then collecting their feces, an activity that would be difficult to achieve. While this limits what conclusions can be drawn on patterns of community ecology, it nonetheless presents empirical evidence of the existence of an organism at a particular place and time, information that is critical to establishing baseline information on species distributions and associations with small mammals.

Many of the species of small mammals captured in this study are known to be omnivorous foragers or herbivores (Table 2) and the presence of a large diversity of fungi contributes to a picture of generalist and opportunistic feeding patterns among these small mammals. Many of these fungi do not produce large, fleshy structures that are likely to be intentionally ingested by small mammals, but their presence may be explained by their symbiotic association with the plants consumed by the animals. For example, *Pleosporales* (specifically *Mycosphaerella*) is a ubiquitous group of plant pathogens with thousands of species (Cheewangkoon et al. 2008; Crous, Braun, and Groenewald 2007) and it is likely that any plant or grain consumed would contain members of this group. Similarly, we found 409 ASVs of *Mucorales*, a nearly cosmopolitan order of mostly saprobic soil- and dung-dwelling fungi, but containing diverse ecologies including animal pathogenic, plant endophytic, and coprophilous lifestyles (Benny, Humber, and Voigt 2014; Walther, Wagner, and Kurzai 2019; Grit Walther, Wagner, and Kurzai 2019; G. Walther et al. 2013). Omnivorous rodent dung is the most common source for a variety of Mucorales and related fungi (Benjamin 1959). Due to the methodology we used for recovering feces (extraction from the intestinal tract during specimen processing), there would have been an extremely limited opportunity for Mucorales fungi to colonize them from the environment (we also did not detect them in our negative controls, further contradicting an environmental contamination hypothesis). This strongly suggests that these fungi were present in the feces when they were sampled. However, it is difficult to ascertain if *Mucorales* fungi may persist within the gut of small mammals or were consumed, possibly indirectly as symbionts of plants consumed by the animals or during coprophagy, a common diet characteristic of many small mammals (Langer 2002).

Of the many fungi detected, 373 ASVs were putative macrofungi that were potentially ingested intentionally by the animals. The presence of a diversity of *Agaricomycotina* (273 ASVs) (the main mushroom-forming fungi), as well as truffle-forming fungi in *Pezizales* (100 ASVs), suggests a tendency toward the consumption of macro- and hypogeous sporocarps common to forested areas. However, no species of fungus was found exclusively in a species or lineage of mammals, nor was there any difference in composition of fungi detected between sexes within a species (Supplementary figure 3). Taken together, our results indicate that small mammals we surveyed are largely opportunistic and generalist fungus feeders that collectively contribute to fungal dispersal. The degree to which these mammals contribute to the dispersal of any given fungal species is unclear. However, the most common and diverse macrofungal genus was the hypogeous *Rhizopogon*, found in 65 of 138 samples (47%). The genus *Rhizopogon* forms hypogeous (belowground) sporocarps that are otherwise difficult to survey by traditional methods, indicating a potentially large role that active foraging by a diversity of small mammals may play in the dispersal of these fungi.

The preponderance of *Rhizopogon* spp. in our samples provided an opportunity to investigate patterns of specialization. Due to its hypogeous reproductive morphology and cosmopolitan distribution, as well as its importance to forest ecology and small mammal diet (Stephens and Rowe 2020), we chose to focus on *Rhizopogon* as a model for determining how these data can contribute to establishing known distributions of fungal species and their impact on formal conservation assessment. However, our ITS2 barcode sequencing targeted the total fungal community, not just *Rhizopogon*, and could therefore be used in the future to address larger-scale questions as well as to apply our methods to a number of taxa that are difficult to locate or rely on animals for dispersal. *Rhizopogon* is a well-known, globally distributed ectomycorrhizal symbiont of the conifer tree family *Pinaceae*, with most species specializing on a genus or subgenus (Cairney and Chambers 1999). *Rhizopogon* spp. form dense and persistent spore banks in soil (Grubisha, Bergemann, and Bruns 2007) Wildfires are a common disturbance in many forest ecosystems, and pine trees (genus *Pinus)* in particular have a fire-adapted ecology and are ecologically dominant in many fire-prone ecosystems (Badik, Jahner, and Wilson 2018). Some *Rhizopogon* species have been shown to have a competitive advantage in host root tip colonization following conditions that mimic wildfire disturbance (Izzo, Canright, and Bruns 2006), indicating that they are important to pine stand reestablishment following a fire (Baar et al. 1999). The ectomycorrhizal habit, specialization on fire-adapted genus *Pinus*, and success in forming mycorrhizae following conditions approximating fire disturbance make *Rhizopogon* a critical community member in the overall health of forest ecosystems and therefore establishing a baseline for their association with the small mammals that serve as their primary dispersal vectors is important for understanding community assembly and habitat recovery following disturbances such as fire.

Through DADA2 we were able to identify 11 species of *Rhizopogon* that could be assigned to known taxa. However phylogenetic analysis was necessary to fully characterize the total *Rhizopogon* diversity. After clustering our unassigned ASVs we produced 11 additional unassignable but distinct *Rhizopogon* OTUs for analysis. However, we were unable to assign them to any known species hypothesis with phylogenetic analysis. This suggests that we may have detected undescribed *Rhizopogon* spp., or that the reference database is incomplete and inadequate for the task, a problem that is becoming increasingly more problematic (Hofstetter et al. 2019). When comparing our *Rhizopogon* phylogenetic diversity (of 22 informative species ASVs) we found no discernible pattern of host diet specialization and even found that variation occurred between samples of the same species collected in close proximity to one another (supplementary table 1). These findings suggest an opportunistic rather than targeted feeding pattern, which has been well characterized in a number of small mammal species (Stephens and Rowe 2020).

The small mammals we used as proxies for the fungi we detected in their feces extended the known distribution of several *Rhizopogon* spp. and were consistent with the current understanding of *Rhizopogon* ecology and phenology. While imperfect, using small mammals as a proxy allows for coarse-scale identification and sampling that can be done quickly and relatively inexpensively without relying on the labor-intensive, repeated field surveys over long periods of time that characterize traditional systematic fungal community surveys. Using bioclimatic data based on the geolocation for each sample containing *Rhizopogon*, we were able to associate over 70% of our sample variability to seasonality, precipitation, and temperature. Additionally, we found that samples tended to cluster more closely together based on the season in which they were collected. While we did not find any striking trends to suggest new insights to the ecology of *Rhizopogon,* instead we were able to discern trends that were consistent with those already known from decades of previous studies, providing external validation for the ecological relevance of our method. Further, we were successful in being able to expand our knowledge of *Rhizopogon* diversity and distribution within the arid western United States, of which little data were previously collected (Supplementary Figure 1).

One practical application of our results is the impact of the expansion of known ranges on conservation criteria such as the IUCN Red List’s Extent of Occurrence (EOO) and Area of Occupancy (AOO). These measurements work best with a large number of data points to compare, but due to the inherent difficulties of finding and documenting fungi (especially over long periods of time), the measurements most likely under-represent the full extent of most fungi. Utilizing Geocat, a tool specifically created to use geospatial information to aid in the assessment of endangered species proposals, ten of the 11 *Rhizopogon* species assigned to known taxa could be considered Endangered according to their AOO. Generally, the addition of fecally-derived geopoints helped to extend the EOO and AOO for all *Rhizopogon* species detected, except in the case of *R. roseolus* and *R. occidentalis*, which decreased. Based on their AOOs, ten of the 11 species we identified would be considered Threatened under IUCN guidelines, with *R. guzmanii* being considered Critically Endangered. In addition, utilizing the presence or absence of a *Rhizopogon* spp., we were able to account for 65 new occurrences in a single year, compared to an average of 7.2 per year across the MycoPortal database (Supplementary Table 2). Realistically, we believe that our data do not point to *Rhizopogon* being highly endangered, but rather to a designation of data deficiency (DD) (Rodríguez et al. 2015). These organisms are often difficult to find, ephemeral, and even harder to track year to year, meaning that more focus needs to be placed on projects dedicated to understanding the population and distribution over time if before we can even begin to understand loss of diversity in this group.

However, many of these species have extremely small occurrence numbers, such as in the most extreme case of *Rhizopogon guzmanii* (n=4, supplementary figure 2), with many having fewer than 30 identifications in the last 50 years (Supplementary Table 2) . For comparison, the Giant Panda, the world’s most iconic conservation project, had 663 animals in captivity in the year 2021 (https://wwf.ca/species/giant-pandas/). This would suggest that if conservation efforts were evenly spread out, some species of *Rhizopogon* should be receiving 150x the funding Giant Pandas receive due to raw numbers, alone. The bulk of conservation effort today is placed on large, culturally significant organisms, which are often plants and animals (Cao, Wu, and Yu 2021; Gonçalves et al. 2021). Relative to these more conspicuous groups, fungi are severely understudied and we run the risk of losing species before we even know they exist or understand their ecological roles. Given the enormous ecological importance of fungi, this oversight has the potential for devastating effects to the ecological health of our planet. OUr study presents one way in which we can increase the rate at which fungi can be documented and to elucidate interactions of other organisms with *Fungi* that may be critical to the health of threatened populations or species. By improving our knowledge about what fungal taxa can be found and where we can better focus conservation efforts and allocate funding in an impactful and targeted manner to help preserve fungal diversity and achieve a more comprehensive view of ecosystem health.

## Methods

### Field methods and mammal specimen preparation

Fecal samples were obtained from small mammals (shrews, rodents, and lagomorphs < 500 g weight) collected during the course of field surveys conducted to determine patterns of local species richness and abundance. Small mammals were collected by removal trapping using Victor and Museum Special lethal snap-traps (Woodstream Corp., Lancaster, PA) baited with a mixture of peanut butter and rolled oats, or Sherman live traps (H. B. Sherman Traps, Inc., Tallahassee, FL) baited with mixed scratch grains. At each sampling locality, traps were set in multiple discrete traplines (of 10–50 traps) across the full range of available microhabitats. Traps were spaced 3–5 m apart and placed in runways, beside fallen logs, by burrow openings, under available cover, or in other locations with probable small mammal activity. Live-trapped animals were humanely euthanized with isoflurane (MWI Animal Health, Boise, ID). Fecal samples were also obtained from fresh road-killed animals (larger rodents and lagomorphs) salvaged opportunistically. Small mammal collecting was done under permit from state wildlife agencies and the US Fish and Wildlife Service. Field methods followed guidelines of the American Society of Mammalogists (Sikes and the Animal Care and Use Committee of the American Society of Mammalogists 2016) and were approved by the Institutional Animal Care and Use Committee of the University of Utah (protocol #s 18-01007 and 18-01008).

Captured and salvaged animals were preserved as museum voucher specimens. Some specimens were prepared in the field, but most animals were frozen with dry ice in the field and moved to a freezer for later preparation in the laboratory. Prior to the preservation, standard external measurements were taken including total length, length of tail vertebrae, length of ear, length of the hindfoot (in mm), and body weight (in grams), and a liver tissue sample was preserved in 70% ethanol. Fecal samples were taken directly from the intestinal tract, placed in labeled Eppendorf tubes, and frozen. Identifications of voucher specimens were verified following final preparation. Taxonomic nomenclature for mammals follows (Solari and Baker 2007)

### DNA Isolation and Sequencing

Fecal pellets were processed using the Zymo Research DNA miniprep kit (#D4300, Zymo Research, Irvine, CA) following the protocol for fecal samples. Feces were processed by placing either a single full pellet or a subsection of a larger pellet to meet the input requirements suggested by the manufacturer (200mg max). Pellets or pellet fragments were homogenized by placing them in 2.0 mL screw-cap tubes containing Zymo lysis solution and beads then and shaking them in a BeadBugTM microtube homogenizer (#Z763713, Sigma) for 300 seconds at speed setting 400.

Genomic DNA was then amplified and sequenced using a PCR-based dual index strategy to allow for samples to be pooled and sequenced using high throughput Illumina technology. To enable efficient multiplexing of each sample, we employed a two-step amplicon protocol that first amplifies the marker gene using primers that include Nextera adapter tails, then using this as a template for a second round of PCR to add unique indices and Illumina flow cell adapters (Gohl et al. 2016). First, the internal transcribed spacer 2 (“ITS2”) region of the ribosomal RNA cistron was PCR-amplified using the primer pairs 5.8S-Fun Nextera and ITS4-Fun Nextera, (Taylor et al. 2016) with Illumina Nextera adapter tails. Sequencing libraries were prepared from these amplicons by using a 1:99 dilution as a template for a second round of PCR with indexing primers consisting of Nextera adapter, unique 8-bp index, and Illumina flow cell adapter. The resulting DNA was then cleaned and normalized across all samples using the AxyPrep MagTM PCR Normalizer Protocol (Axygen Biosciences, Union City, CA). Normalized sample DNAs along with two negative controls were pooled and submitted for Illumina miSeq 2 x 250 PE sequencing at the University of Utah Genomics Core facility. Of the two Negative controls, one had faint amplified sequences determined by gel electrophoresis while the second did not. However, each control did not generate any identifiable sequence and was completely filtered out during demultiplexing and DADA2 quality filtering.

### Taxonomic assignment of sample contents and Data visualization

The DADA2 pipeline (Callahan et al. 2016) was used to trim and error correct raw sequencing reads, and to generate a table of amplicon sequence variants (ASVs) (Callahan, McMurdie, and Holmes 2017) with default parameters. Each ASV was assigned taxonomically based on similarity to species hypothesis sequences in the UNITE general release database version 8.2 (04 April 2020) of the fungal taxon (Abarenkov et al. 2010).

After taxonomic Identification by DADA2, ASV data was combined with metadata for each small mammal fecal sample and analyzed using the R package Phyloseq version 1.32.0 (McMurdie and Holmes 2013). Pre-processing of the data set included removal of any samples that exhibited only 1 ASV or less. Phyloseq was then used to analyze and visualize multiple aspects of our data such as alpha diversity (plot_richness function) measurements. Further, a total community analysis was done using the package Metacoder version 0.3.4 (Foster, Sharpton, and Gr 2017) to illustrate the total community composition.

### Phylogenetic analysis of non-assignable species and faith’s phylogenetic diversity calculations

Samples containing sequences corresponding to the *Rhizopogon* genus were subsampled from the total community for further analysis. Sequences that were not identified to the species rank by DADA2 were clustered together into OTUs based on a 99% similarity cutoff using Vsearch version 2.15.1 (Rognes et al. 2016) and then subjected to phylogenetic analysis to further clarify phylogenetic and diversity relationships.

Data sets for phylogenetic analysis were created by these OTU’s, all species hypothesis reference sequences as defined by the UNITE database for the Family *Rhizophoraceae* in combination with species identified *Rhizopogon* ASVs. All sequences were run through ITSx version 1.1.2 (Bengtsson-Palme et al. 2013) to normalize each sequence to ITS2, then Multiple sequence alignments from these datasets were generated using the L-INS-i algorithm in MAFFTv7 (Katoh 2002) and phylogenetic trees were inferred under maximum likelihood using IQ-TREE with the model of molecular evolution automatically determined and branch support estimated by ultrafast bootstrapping (Nguyen et al. 2015; Minh, Nguyen, and von Haeseler 2013). Finally, community data for each tree was transformed to presence/absence reporting and combined with their respective phylogenetic tree to measure faith’s phylogenetic diversity using the Picante R package version 1.8.2 (Kembel et al. 2010).

### Principal component analysis and Bioclimatic variable analysis

Bioclimatic variables were downloaded from the worldclim database (Fick and Hijmans 2017), based on the geolocation recorded from each collected small mammal sample which included sequences assigned to the *Rhizopogon* genus. Principal component analysis was performed using the PCAshiny function of the R package Factoshiny version 2.4 (Lê, Josse, and Husson 2008). Factoshiny is a GUI implementation of the FactoMineR package. Visualization was generated by altering the components to only display in dimensions 1 and 2 (corresponding to 77.32% of variability) and to only label the top 5 contributing environmental variables. Individual PCA was conducted on the individual samples and colored by the quantitative sample values of sex and season collected respectively with confidence ellipses added.

### Analysis of Extent of Occurrence and area of occupancy for *Rhizopogon* species

*Fungi* function as the main decomposers of organic material, form-critical symbiotic partnerships affecting plant health, and serve as a nutritional food source for animals. Therefore, a thorough understanding of fungal diversity and distribution is extremely important to identify targets under threat of extinction and to develop appropriate methods for their conservation. As proof of concept, we evaluated whether our data and analyses can assist in making meaningful conclusions about the distribution of *Rhizopogon,* and be used to inform possible conservation efforts. Information needed to accurately inform how conservation efforts should be focused is often difficult to ascertain and highly complex (Vane-Wright, Humphries, and Williams 1991; Callmander et al. 2007; Laity et al. 2015; Juffe-Bignoli et al. 2016). Baseline data to establish the currently known extent of occurrence and area of occupancy of specific *Rhizopogon* species was created by using the geospatial analysis online tool GeoCat (Bachman et al. 2011) at <u>GeoCAT</u>, using publicly available observation and collection data from from the global biodiversity informational facility (GBIF) and MycoPoral respectively. Geospatial coordinates were generated by subsetting our data set for samples that had ASVs corresponding to each *Rhizopogon* species (supplementary table 3). In each sample, all ASVs assigned to a specific *Rhizopogon* species were treated as a single occurrence and geospatial coordinates of the corresponding trapped small mammal were used as a proxy for that specific *Rhizopogon* species presence within the environment.

## Supporting information

All Tables in xlsx format

## Data Availability

Raw short-read sequences for this project have been deposited in SRA through the bioproject PRJNA764247. Raw tree files, Phyloseq, and Geocat objects have been deposited in Figshare under https://doi.org/10.6084/m9.figshare.15105462.v1. Any code or specific script requests should be sent to the corresponding author.

## Author contributions

Alexander J Bradshaw processed samples, generated and conducted protocols to sequence DNA, coding to process and analyze data, in addition to preparation and writing of this manuscript. Kendra Autumn participated in processing specimen samples as well as providing assistance in data compilation and analysis, artistic figure production, and the preparation and writing of this manuscript. Eric Rickart performed all fieldwork and small mammal dissection and fecal sampling as well as providing critical insight into the ecology of the specimens used in this study. Bryn T.M. Dentinger provided funding, experimental guidance, and assisted in the preparation and editing of this manuscript.

## Acknowledgments

We acknowledge the Natural History Museum of Utah for their commitment to collaborative science as well as the Genomics Core Facility, a part of the Health Sciences Cores at the University of Utah for their input and high-quality work. We are grateful to Shannen Robson for curatorial support. Further, we wish to acknowledge the dedicated and hard work performed by Allyson Jelito, Bryce Alex, and Isabelle Galland to help process many of these samples for DNA sequencing.

## Supplementary Figures and Tables

**S1: Geopoints comparison of our samples and mycoportal collection geopoints**

**S2: expansion of EOO for *R. guzmanii***

**S3: Principal component analysis with confidence ellipse of *Rhizopogon* presence based on sample sex**

**S4: Alpha diversity measurements (shannon and choa1) of all *Rhizopogon* species in each sample**

**ST1: Faiths PD and species richness of all *Rhizopogon* containing samples**

**ST2: Comparison of public mycoportal database *Rhizopogon* Identifications to our study over time**

**ST3:Table of all samples containing *Rhizopogon* ASVs**

