## Supplemental Figures, S1-S4 for "On the Origin of Feces: Fungal diversity, distribution, and conservation implications from feces of small mammals"

Supplementary figures:

S1:Overall collection point distribution

Depicts geolocation as well as the total count for each state sampled. In red are mycoportal collections, shown mostly around major cities in the pacific northwest.

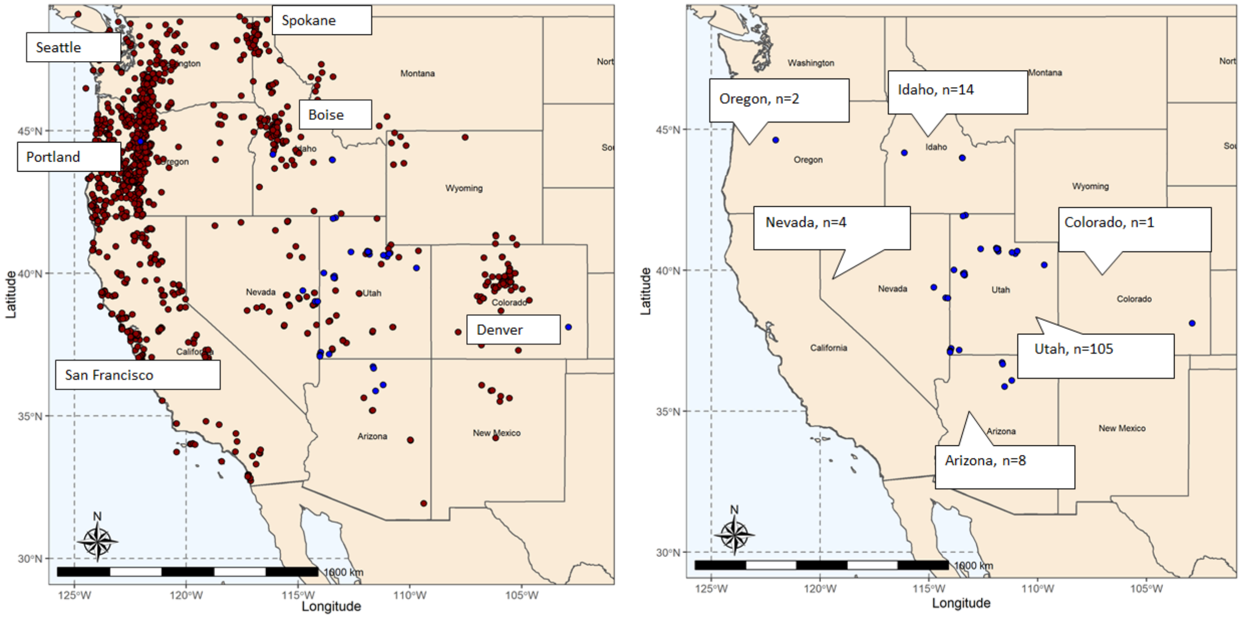

S2: Extending the Extent of Occurrence for *Rhizopogon guzmanii*

Left: Known EOO of *Rhizopogon guzmanii* as determined by four records pulled from GBIF.

Right: Known EOO as well as Collection point which was found to contain *Rhizopogon guzmanii* ITS sequences, extending the possible known range to include both central Mexico as well as northern Utah.

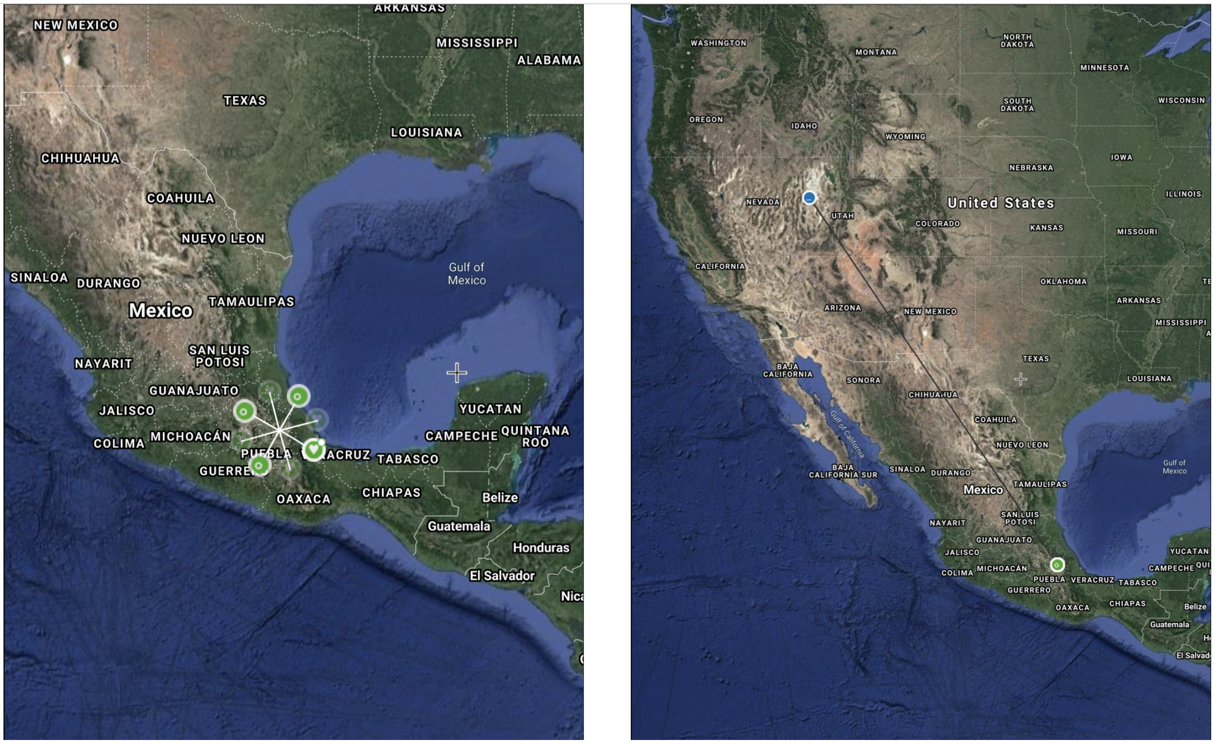

S3: PCOA for sex of small mammals

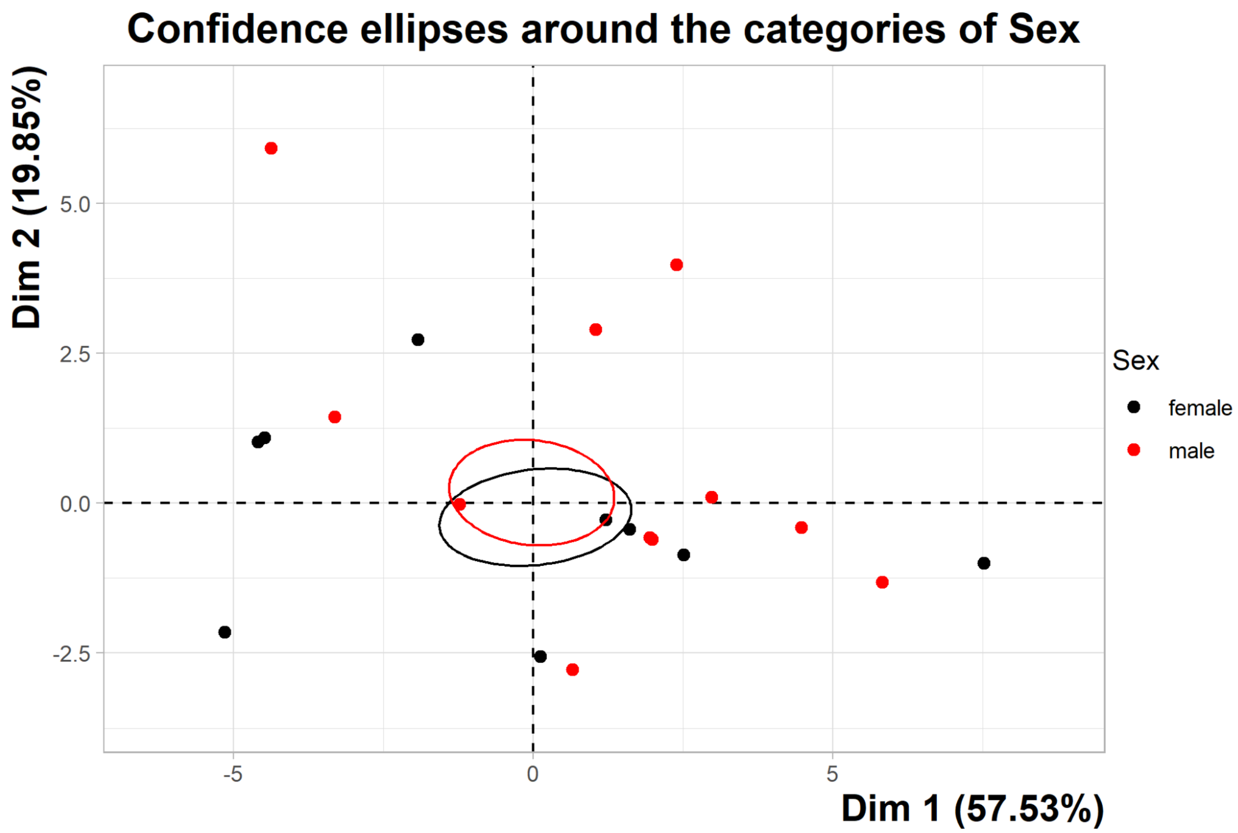

### S4:Alpha Diversity of *Rhizopogon* containing samples

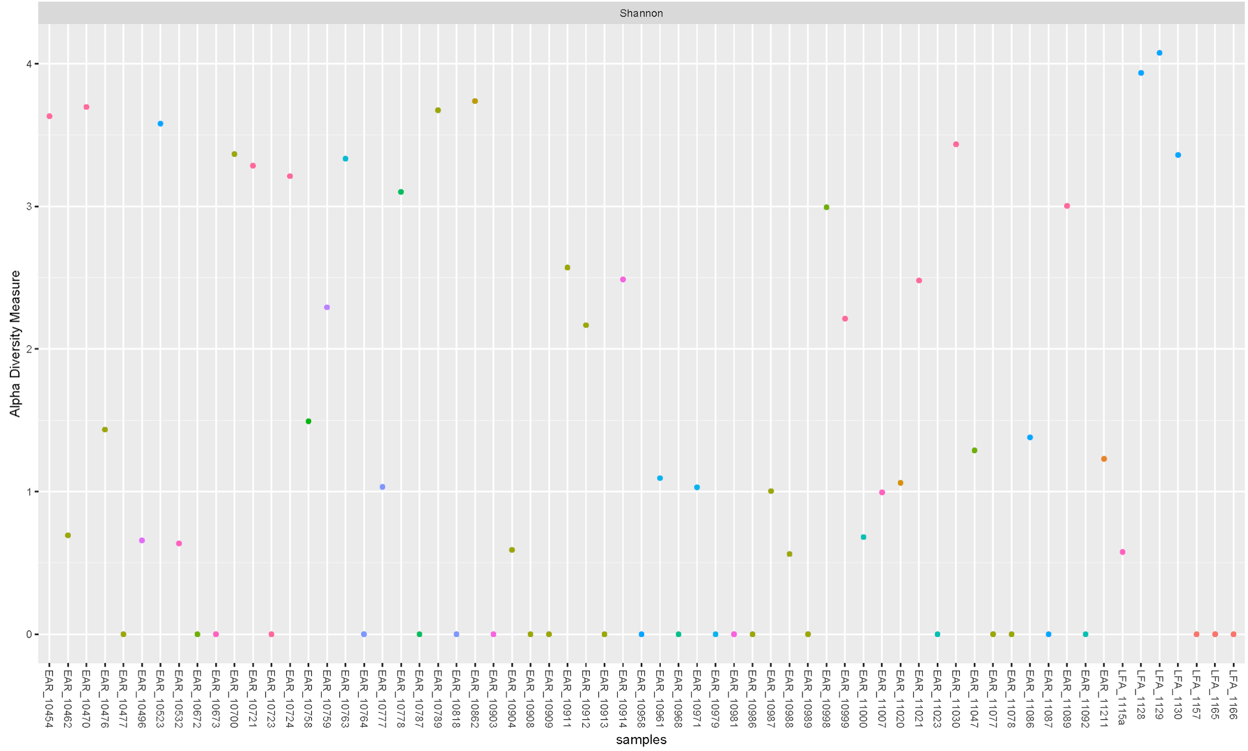


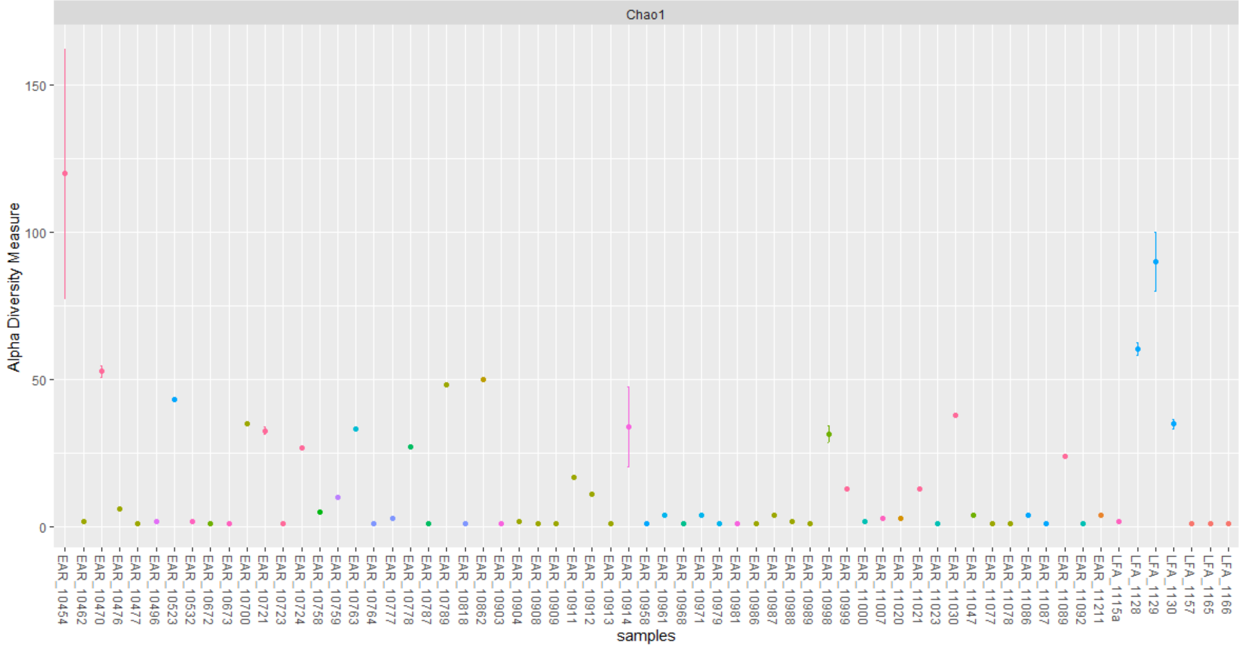
